# Noncoding RNA’s competing endogenous gene pair as motif in serous ovarian cancer

**DOI:** 10.1101/2022.04.04.486923

**Authors:** Lixin Cheng, Xubin Zheng, Ning Zhang, Jing Gao, Kwong-Sak Leung, Man-Hon Wong, Shu Yang, Yakun Liu, Ming Dong, Huimin Bai, Lin Kang, Haili Li

## Abstract

Understanding the regulatory mechanisms in serous ovarian carcinoma (SOC) is critical for its diagnosis and targeted therapy. However, some critical motifs in the competing endogenous RNA (ceRNA) network in SOC were still undiscovered. We profiled a whole transcriptome of eight human SOCs and eight controls and constructed a ceRNA network including mRNAs, lncRNAs, and circRNAs. We hypothesized the noncoding RNA’s competing endogenous gene pairs (ceGPs) relationship for the mRNA–ncRNA–mRNA motifs in the ceRNA network. Then, we proposed the denoised individualized pair analysis of gene expression (deiPAGE) to identify mRNA–ncRNA–mRNA motifs from integrated multi-cohorts. 18 cricRNA’s ceGPs (cceGPs) were identified and fused as an indicator (SOC index) for SOC discrimination, which carried a high predictive capacity in independent cohorts. The index was negatively correlated with the CD8+/CD4+ ratio in tumour-infiltration, reflecting the migration and growth of tumour cells in ovarian cancer progression.

## 1. INTRODUCTION

Ovarian cancer is the most lethal malignancy worldwide in gynaecology [1]. According to estimates from the American Cancer Society, 1 in 78 women will suffer from ovarian cancer. Furthermore, around 21,750 will be newly diagnosed and 13,940 will die from ovarian cancer in 2020 [2]. Epithelial ovarian carcinomas account for 90% of ovarian cancer cases and serous ovarian carcinoma (SOC) is thus far the most common subtype in epithelial ovarian carcinomas [3, 4]. Understanding the regulatory mechanisms in SOC is critical for the diagnosis and the treatment of SOC.

Competing endogenous RNAs (ceRNAs) represent a regulation mode in which ceRNAs interact with other RNAs by competing for the shared target microRNAs [5, 6]. Chiu et al. validated the ceRNA regulatory network in prostate and breast adenocarcinomas [7]. Liang et al. constructed a ceRNA regulatory network for mesenchymal ovarian cancer and identified the downregulation of lncRNA pro-transition associated R (PTAR) potentially inhibiting cancer metastasis by sponging miR-101 [8]. PTAF is a pivotal regulator of the epithelial-to-mesenchymal transition promoting the invasion–metastasis cascade of ovarian cancer. Furthermore, they demonstrated that the overexpression of PTAF can upregulate snail family zinc finger 2 (SNAI2) by directly sponging miR-25, leading to the promotion of ovarian cancer epithelial-to-mesenchymal transition and invasion [8]. Wang et al. found that lncRNA small nucleolar RNA host gene 16 (SNHG16), acting as a ceRNA, played important roles in the immune processes and was upregulated in myasthenia gravis patients [9].

Nevertheless, investigations about ceRNAs mainly concentrated on the regulatory relation between individual ncRNAs and mRNAs, i.e., ncRNA – miRNA – mRNA. Studies about the regulatory motifs in the ceRNA network did not move forward, such as extending the regulatory chain or combining with other regulatory nodes. Moreover, current research on ncRNA in the ceRNA network in ovarian cancer merely focused on lncRNAs [10–14], given that no sufficient high-throughput circular RNA (circRNA) expression data are publicly available for ovarian cancer. As a layer of the gene regulatory network, circRNAs feature a variety of biological processes, including tumour cell proliferation, migration, and invasion [15]. Identifying motifs from ceRNA and investigating the correlation between circRNAs and other types of RNAs will shed light on the underlying molecular mechanisms of ovarian cancer.

In this study, we hypothesized the triangular relationship of ncRNA’s competing endogenous gene pair (ceGP), where the genes and ncRNA composed a mRNA – miRNA - ncRNA – miRNA – mRNA motif in the ceRNA network and the two genes reversed in their expression between SOC and normal controls (**Figure 1A**). First, we generated expression profiles from eight SOCs and eight normal ovary samples to characterise the full RNA expression pattern of human SOC using Agilent microarrays (**Figure 1B**). To comprehensively understand the alteration of RNAs, we assessed the differentially expressed mRNAs, long non-coding RNAs (lncRNAs), and circular RNAs (circRNAs), and constructed a ceRNA network based on them. Next, we proposed the denoised individualized pair analysis of gene expression (deiPAGE) to select ncRNA’s ceGPs (nceGPs) and developed a diagnostic indicator (SOC index) for SOC discrimination based on three cohorts including the self-profiled one. Validation on two independent cohorts and one blood cohort were performed. After that, we investigated the correlation between the SOC index and tumour infiltrating cells including the two typical T cells, CD4+ and CD8+. Functional enrichment and literature study were also carried out for the gene pairs in SOC index. Finally, a cricRNA’s ceGP (cceGP) *BBS4-circHUNK-PRC1* was illustrated as an example for the nceGP relationship in SOC.

**Figure 1.**
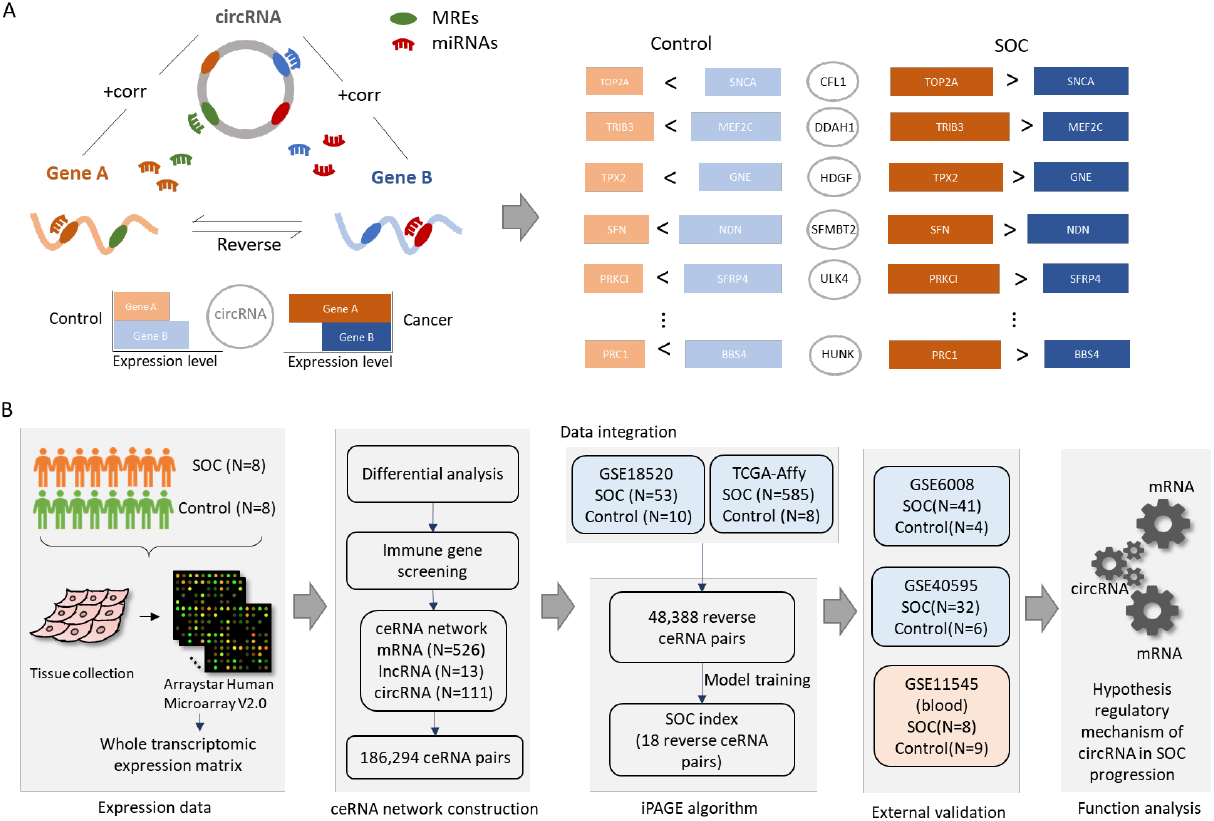
Overview of this study. A) Illustration of circular RNA’s competitive endogenous gene pair (cceGP), where the two genes reversed in their expression between serous ovarian carcinomas (SOCs) and normal controls. Multiple pairs were identified to discriminate SOC from normal controls. B) The workflow to study cceGPs as signature in SOC.

## 2. MATERIALS AND METHODS

### 2.1 Patients and samples

16 patient samples were included in this study (**Table 1**). Eight patients with ovarian cancer and eight patients with cervical cancer (normal ovarian tissue specimens) were recruited sterilely at the Fourth Hospital of Hebei Medical University between 2015 and 2017. This study was approved by the institutional review board of Hebei Medical University. Both cancerous and normal ovarian tissues were quickly excised and snap-frozen in a liquid nitrogen tank at −80°C until further use.

**Table 1.**
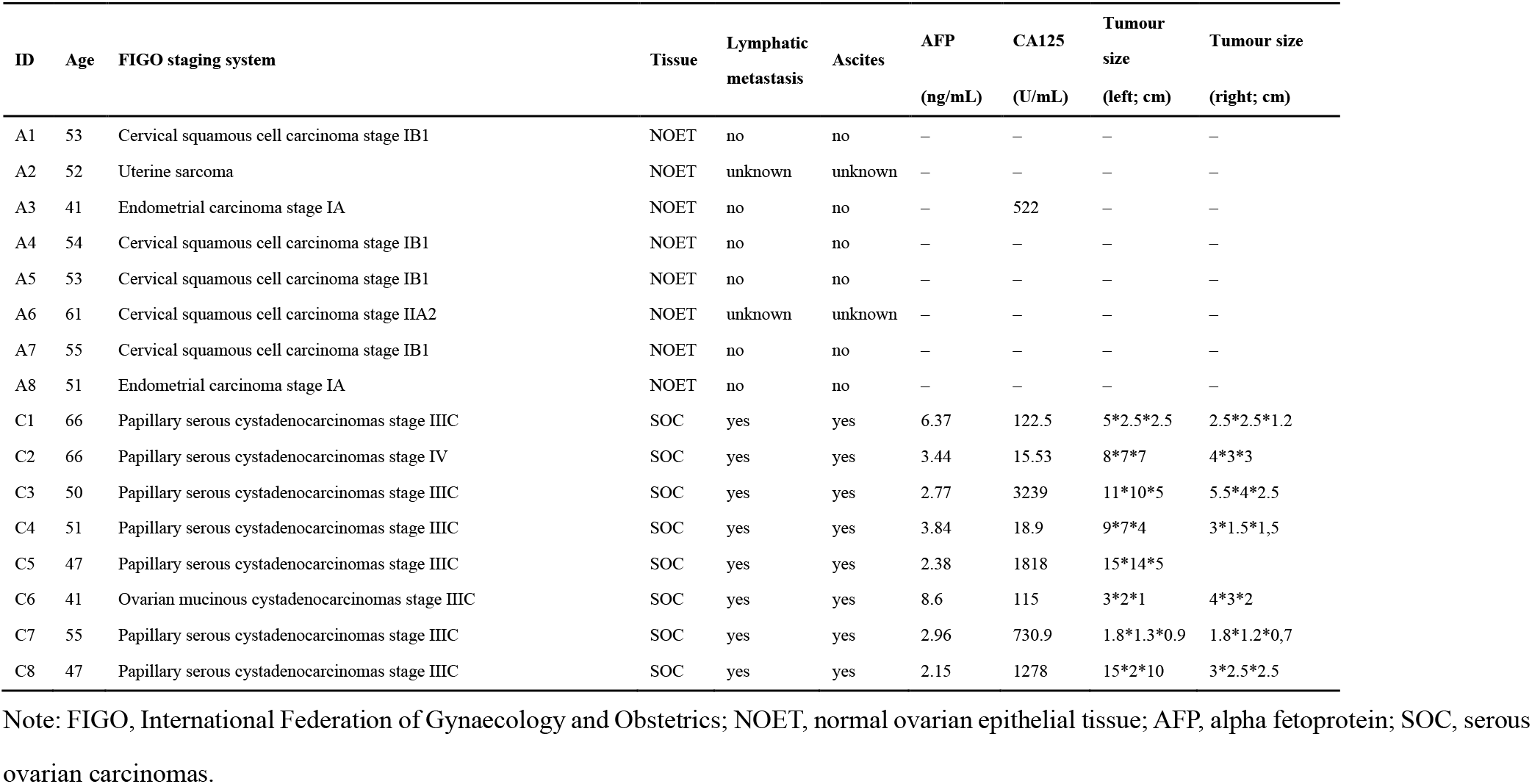
Clinical characteristics of patients with ovarian cancer and control samples.

### 2.2 RNA extraction

Tissues were homogenised in the TRIZOL reagent (Invitrogen, USA) using a Qiagen Tissuelyser. Total RNA was extracted in accordance with the manufacturer’s protocol and then quantified using a NanoDrop ND-1000 spectrophotometer (Thermo Fisher Scientific, Waltham, MA, USA). The RNA integrity of each sample was assessed by denaturing agarose gel electrophoresis.

### 2.3 Microarray experiments

To identify deregulated RNAs associated with SOC patient outcomes, we conducted a microarray study and profiled mRNAs, lncRNAs, and circRNAs, respectively. We performed Arraystar Human LncRNA Microarray V2.0 and Arraystar Human circRNA Array V2.0 analyses on all 16 samples. The expressions of lncRNAs and mRNA were quantified using the first platform, while the expressions of circRNAs were measured using the second. Total RNA from each sample was measured using NanoDrop ND-1000. Sample preparation and microarray hybridisation were performed based on the standard protocols of Arraystar (Agilent Technology, USA). Processing RNA was different between the two platforms. For the lncRNA platform, rRNA was removed from total RNA using the mRNA-ONLY™ Eukaryotic mRNA Isolation Kit (Epicentre Biotechnologies, USA). For the circRNA platform, total RNAs were digested with Rnase R (Epicentre, Inc.) to remove linear RNAs and enrich circular RNAs. Then, the two platforms followed the same steps below.

Each sample was amplified and transcribed into fluorescent cRNA utilising a random priming method (Arraystar Super RNA Labeling Kit; Arraystar). The labelled cRNAs were purified using the RNeasy Mini Kit (Qiagen, Germany). The concentration and specific activity of the labelled cRNAs (pmol Cy3/μg cRNA) were measured using NanoDrop ND-1000. Here, 1 μg of each labelled cRNA was fragmented by adding 5-μl 10 × blocking agent and 1-μl 25 × fragmentation buffer, heating the mixture to 60°C for 30 min, and then adding 25-μl 2 × hybridization buffer to dilute the labelled cRNA. Next, 50-μl hybridisation solution was dispensed into a gasket slide and assembled on the circRNA expression microarray slide. The slides were incubated for 17 hours at 65°C in an Agilent Hybridisation Oven. The hybridised arrays were washed, fixed, and scanned using the Agilent Scanner G2505C. We used the Agilent Feature Extraction software (version 11.0.1.1) to analyse the acquired array images.

### 2.4 Datasets

Other than the self-profiled cohort, we downloaded the gene expression of SOC samples from TCGA and GEO (GSE18520, GSE6008, GSE40595 and GSE11545; **Table 2**). We performed differential and functional analysis and constructed a competing endogenous RNA regulatory network based on the self-profiled cohort. We then combined the self-profiled cohort, GSE18520, and the TCGA cohort retrieved from UCSC Xena including 672 samples as a training set for SOC signature identification (**Figure 1B**). GSE6008 and GSE40595 were applied as two independent validation sets. To verify the signature for non-invasive diagnosis, we adopted a blood cohort GSE11545 as another external validation set.

**Table 2.**
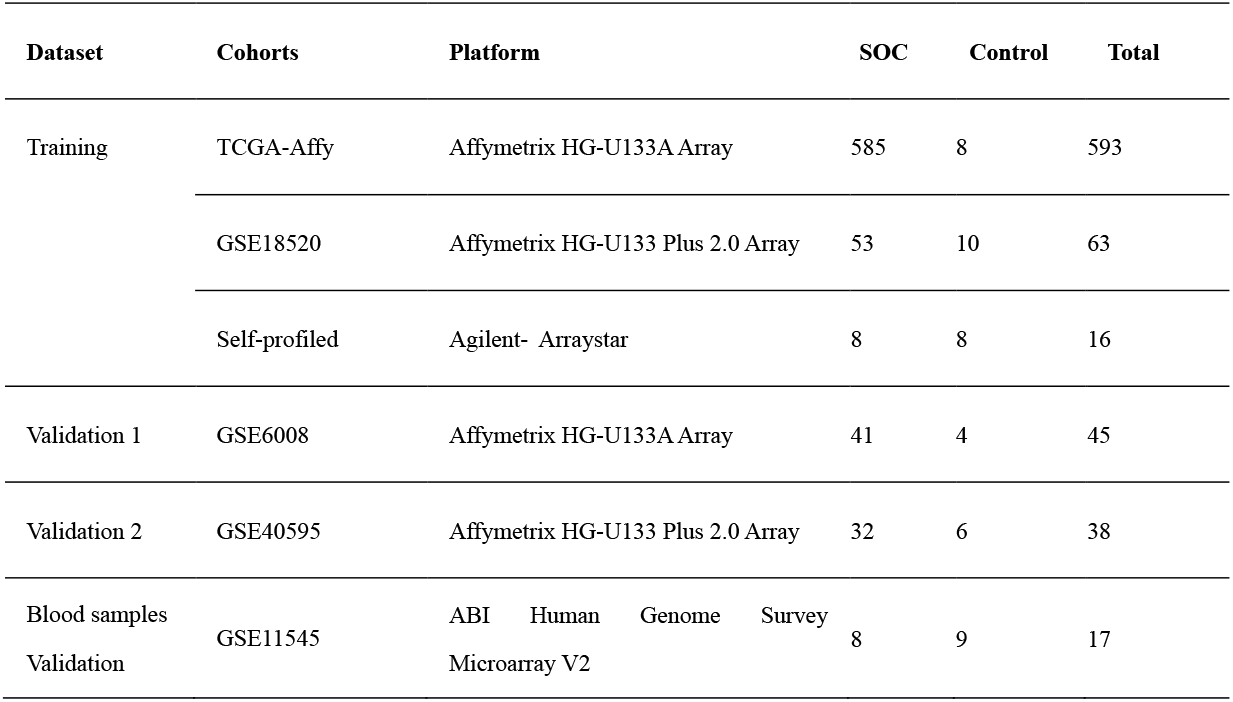
Gene expression datasets used in this study.

| Dataset | Cohorts | Platform | SOC | Control | Total |
| --- | --- | --- | --- | --- | --- |
| Training | TCGA-Affy | Affymetrix HG-U133A Array | 585 | 8 | 593 |
|  | GSE18520 | Affymetrix HG-U133 Plus 2.0 Array | 53 | 10 | 63 |
|  | Self-profiled | Agilent- Arraystar | 8 | 8 | 16 |
| Validation 1 | GSE6008 | Affymetrix HG-U133A Array | 41 | 4 | 45 |
| Validation 2 | GSE40595 | Affymetrix HG-U133 Plus 2.0 Array | 32 | 6 | 38 |
| Blood samples<br>Validation | GSE11545 | ABI Human Genome Survey<br>Microarray V2 | 8 | 9 | 17 |

Gene expressions from the five cohorts were quantified using three platforms, i.e., the Affymetrix HG-U133A Array, the Affymetrix HG-U133 Plus 2.0 Array, and the Agilent-Arraystar. Besides, GSE11545 measured gene expression of SOC in blood using ABI Human Genome Survey Microarray V2.

### 2.5 Differential and functional analysis

The expression profiles of SOC patients and normal controls in the self-profiled cohort were analysed to identify differentially expressed mRNAs, lncRNAs, and circRNAs, respectively. Quantile normalization was used to normalize the profiles to render the data comparable across samples. The significance levels were estimated using the Student’s t-test and, then, adjusted using the Benjamini and Hochberg (BH) multiple testing correction method. The effect size was also taken into account of using the two-fold change [16, 17]. In total, we obtained 1,881 downregulated mRNAs and 1,995 upregulated mRNAs, 1,849 downregulated lncRNAs and 718 upregulated lncRNAs, as well as 122 downregulated circRNAs and 69 upregulated circRNAs.

### 2.6 Competing endogenous RNA network construction

The competing endogenous RNAs (ceRNAs) sponged the same micro RNAs (miRNAs) and the their expression are positively correlated. We used the Pearson’s correlation test to calculate the expression correlation among mRNAs, lncRNAs, and circRNAs in SOC. Then we screened those that shared the same interactive miRNAs through RNAInter [18]. Only RNAs positively correlated (Pearson correlation coefficient (PCC) > 0.5) and interactive with the same miRNAs were considered as ceRNAs for the construction of the competing endogenous regulatory network [5, 19]. The final network was illustrated using Cytoscape [20].

### 2.7 Denoised individualized pair analysis of gene expression

The abundance of genes may vary across different detection platforms or preprocessing methods, but the relative ranking is stable in a pair of genes. Herein, we developed denoised individualized pair analysis of gene expression (deiPAGE). deiPAGE not only considered the relative ranking but also included the effective size between two genes. Moreover, deiPAGE applied the least absolute shrinkage and selection operator (LASSO) for feature selection.

#### 2.7.1 Establishment of ceGPs

The gene expression profiles from different training cohorts were concatenated directly as a training set. Then, we performed pairwise subtraction for all genes to form gene pairs (*g_i_* – *g_j_*) in a single sample (**Figure 2A**). However, the gene expression value may vary due to technical noise and it was denoted as *g* = *g*′ + *ε*, where *g*′ was the true value of the gene expression and *ε* ∈ (−∞, +∞) represented the error of measurement. When subtraction was performed, it became 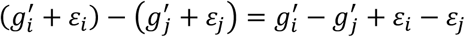. The difference within a pair may not only result from the ground truth of the gene expression 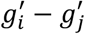, but was also affected by the technical variation *ε_i_* – *ε_j_* when counting RNAs from reads. To remove the relative expression caused by the technical variation, we introduced effective size Δ, where only the difference 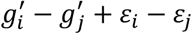 exceeding Δ was regarded as effective. As a result, we applied the function 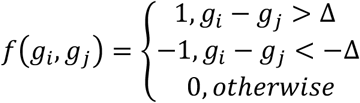 to each gene pair (**Figure 2B**).

**Figure 2.**
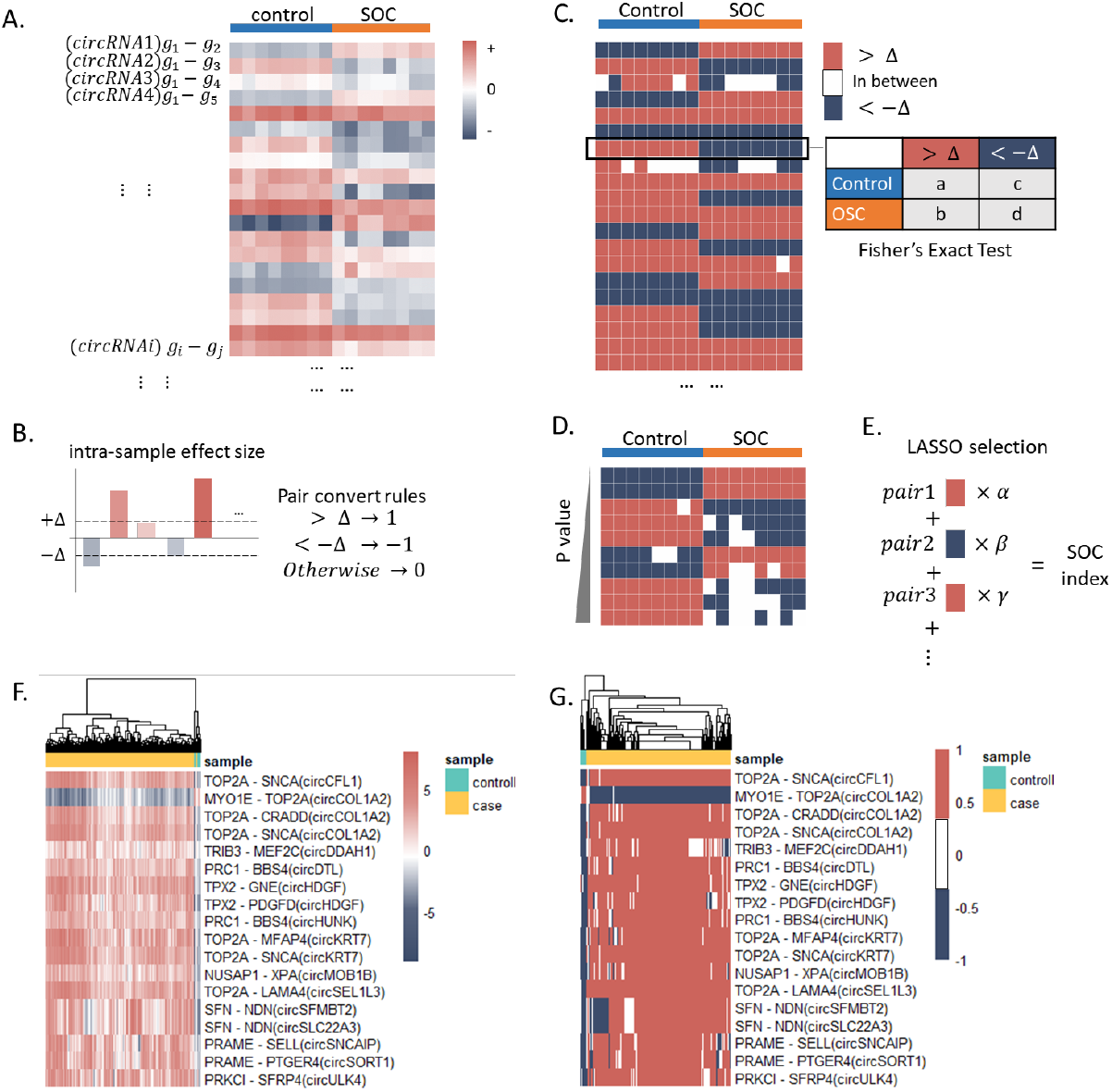
The workflow of denoised individualized pair analysis of gene expression (deiPAGE) to compute the SOC index. A) Subtraction of two genes within each sample of the expression matrix. B) Gene pairs with a large effect size (Δ) were converted to intrasample relative expression indicators (1 or −1). Gene pairs with a small effect size possibly introduced from technical variation were filtered out (0). C) Based on the intrasample rank, a Fisher’s exact test was performed across the population without considering a small effect size for each gene pair. D) Gene pairs were ranked by their P-value and differentially reversed gene pairs (DRPs) were screened. E) Feature selection and construction of SOC index by machine learning method, the least absolute shrinkage and selection operator (LASSO) regression. F) Heatmap showing the subtraction result of the 18 circRNA’s competitive endogenous gene pairs (cceGPs) identified by LASSO. G) Heatmap showing the subtraction result of the 18 cceGPs after conversion.

Following the intrasample analysis, we performed a population analysis to obtain the significance level of the gene pairs. For each gene pair, a contingency table was calculated without considering the samples within the difference threshold, given that those samples are largely affected by noise (**Figure 2C**). We then conducted the Fisher’s exact test, and 48,388 significantly different pairs (False discovery rate corrected *P* < 0.01) were preserved as noncoding RNA’s competing endogenous gene pairs (nceGPs) (**Figure 2D**).

#### 2.7.2 SOC index generation

The least absolute shrinkage and selection operator (LASSO) [21–23] is a regression model with regularization in machine learning, which has been widely used in medical applications [24–26]. It can not only classify different classes, but also prune features to avoid overfitting and improve the generalizability. The linear regression was optimized using the following loss function with an L1 penalty: 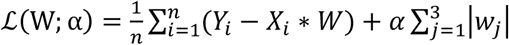, where *n* is the number of samples, *Y* is the label for each sample, *X* is the vector composed by all the nceGPs and the constant term, *W* is the vector of weights for nceGPs, and α is the coefficient for the L1 penalty.

After 48,388 differential gene pairs were obtained from the previous step, resampling was applied to the training dataset due to the unbalanced size of the SOC samples and normal controls, which is a commonly used trick in machine learning to improve model accuracy. LASSO regression model was trained on the resampled training set and 18 circular RNA’s competing endogenous gene pairs (cceGPs) were selected for the final SOC index (**Figure 2E**). The code of deiPAGE algorithm is available at https://github.com/Kimxbzheng/deiPAGE.

### 2.8 Principle component analysis

Principal component analysis (PCA) is a machine learning method to summarize highdimensional data into a few main components for data analysis and visualization. In this study, we performed PCA using sklearn in python to convert the 18 cceGPs into three principal components for visualization of its classification performance (**Figure 4A**).

**Figure 3.**
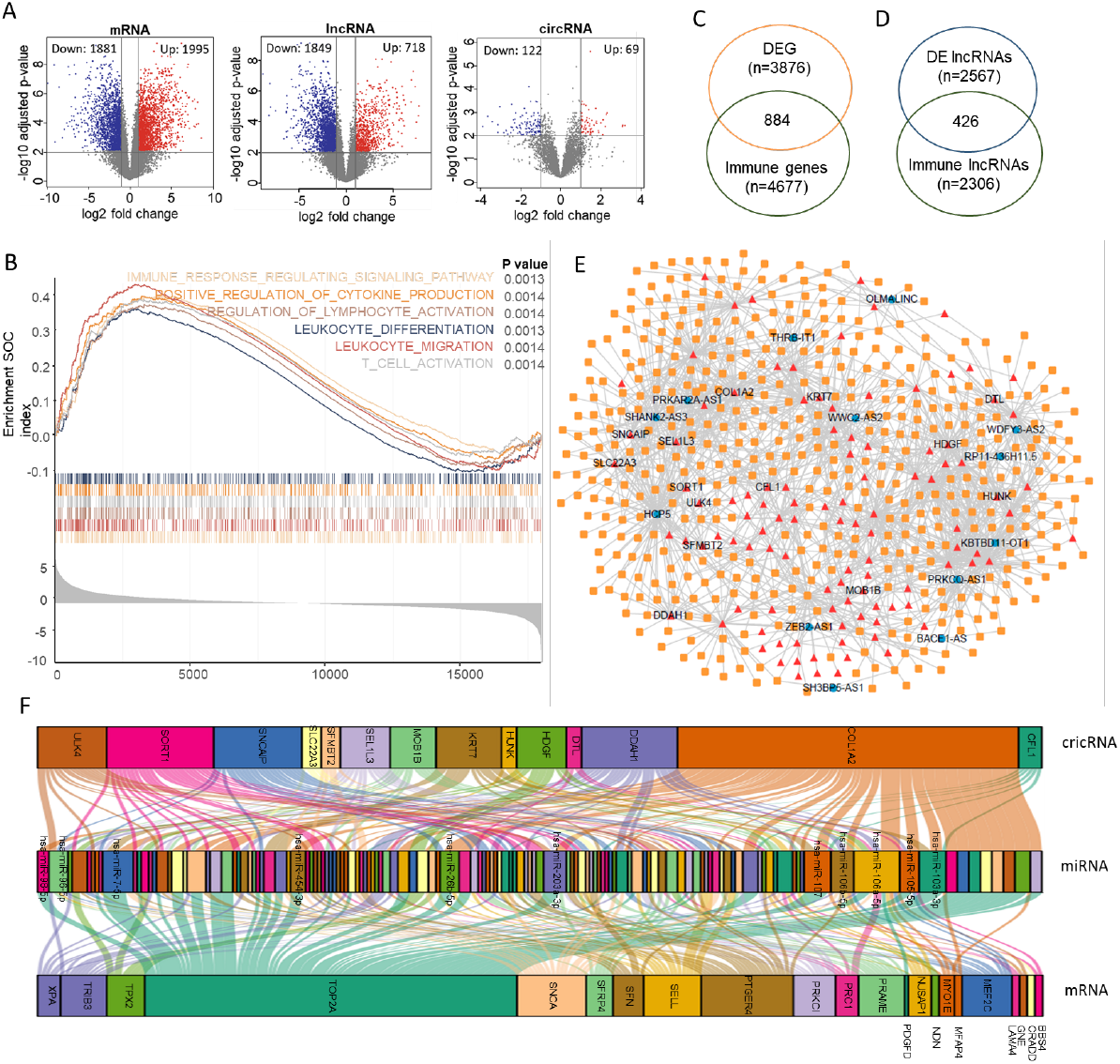
Construction of the competitive endogenous RNA (ceRNA) regulatory network. A) Volcano plot of differentially expressed mRNAs, lncRNAs, and circRNAs. B) Functional analysis of differentially expressed RNAs using GSEA. A majority of the enriched biological processes are associated with immunity. C) Venn diagram of the differentially expressed genes and immune-related genes. D) Venn diagram of the differentially expressed lncRNAs and immune-related lncRNAs. E) The immune-related ceRNA regulatory network composed of mRNAs, lncRNAs, and circRNAs. F) Sankey diagram of the competitive endogenous circRNAs and mRNAs identified by deiPAGE. The miRNAs interacting with them are in the middle layer.

**Figure 4.**
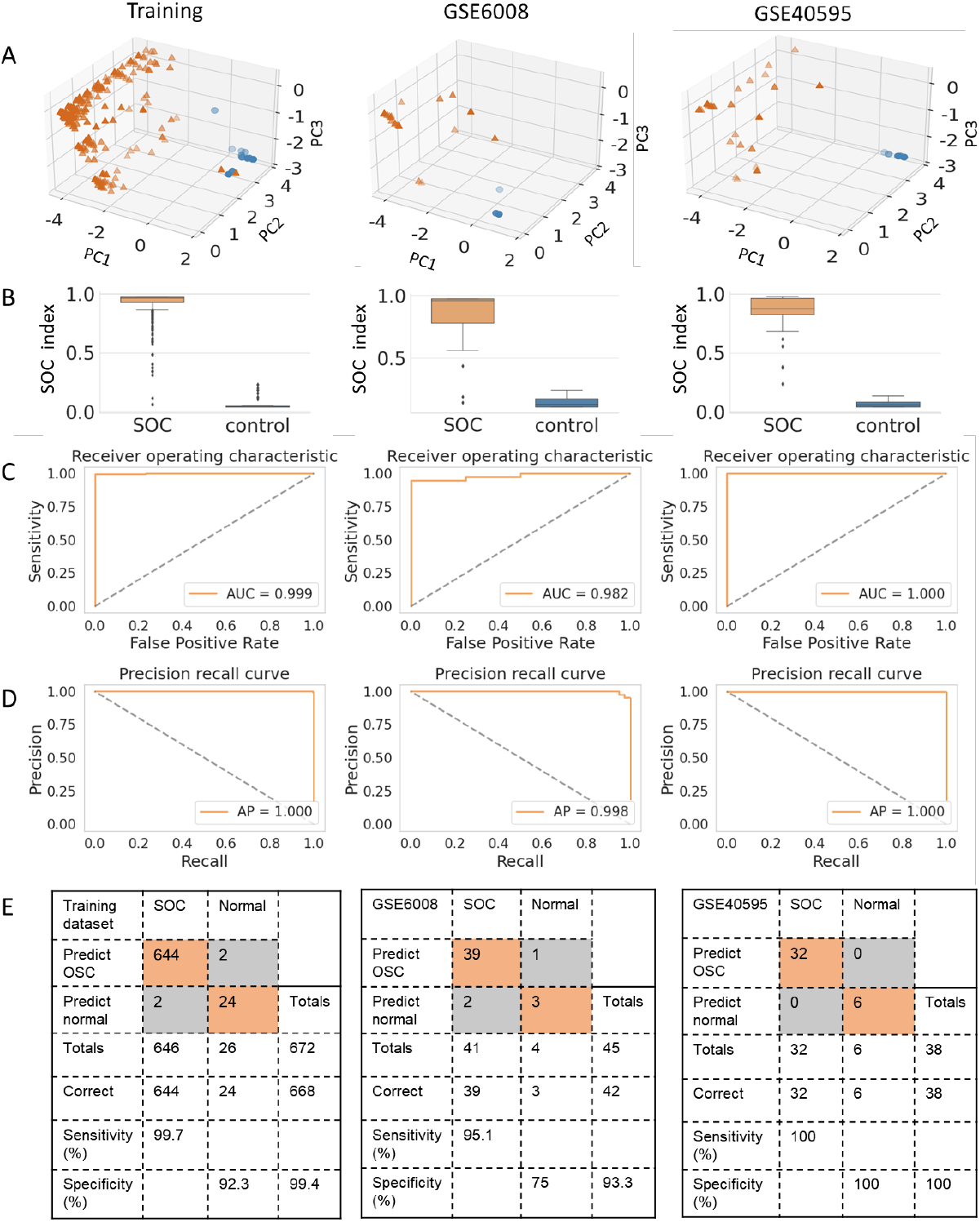
Performance evaluation of the SOC index. A) The three principal components of the 18 cceGPs in the training and validation sets. B) Box plot of the SOC index in the training and validation sets. C) ROC curves of SOC index in the training and validation sets. D) Precision-recall curves of SOC index in the training and validation sets. E) Confusion matrix of SOC index in the training and validation sets.

### 2.9 Correlation analysis with tumour infiltration

To explore the association with immune cell infiltration, Spearman’s rank correlation coefficient was adopted to estimate the correlation between the SOC index and immune cells in cancer (|R| > 0.3 and *P* < 0.01). For each sample, we calculated the levels of immune cell infiltration using the Tumour Immune Estimation Resource (TIMER) [27].

### 2.10 Gene set enrichment analysis and functional enrichment analysis

We downloaded all pathways of Collection 2 (C2) and their gene sets from MSigDB (v7.4) and applied Gene Set Enrichment Analysis (GSEA) [28] with the R package clusterProfiler [29]. The differentially expressed mRNAs were subjected to GSEA [30] for functional enrichment analysis. Hypergeometric test was applied to evaluate the statistical significance of functional enrichment for overrepresented gene sets using the R package clusterProfiler [29]. The 18 cceGPs in SOC index were enriched in Gene Oncology (GO) [31] and Reactome [32].

### 2.11 Statistical analyses

R Project (R x64, version 3.5.2) and Python (version 3.6) were used for statistical computation in this study. Student’s t-test and the Benjamini and Hochberg (BH) multiple testing correction method were used to calculate the differential expression RNAs in the self-profiled cohort. Pearson’s correlation test were applied to calculate the expression correlation among RNAs. Fisher’s exact test and false discovery rate (FDR) correction were employed to compute the significance of gene pair reversal. The LASSO regression was used to screen gene pairs and construct classification model for SOC. PCA was used to visualize the 18 gene pairs. Kaplan-Meier method and log-rank test were used for clinical outcome comparison. Spearman’s rank correlation coefficient was adopted to estimate the correlation between the SOC index and immune cells. Hypergeometric test was applied to evaluate the statistical significance of functional enrichment for overrepresented gene sets. Statistical significance was set at P < 0.01.

## 3. RESULTS

### 3.1 Clinical characteristics and differential analysis

Aged between 41 and 66 years, eight serous ovarian carcinoma (SOC) patients and eight patients without SOC were included in this study (**Table 1**). Patients with SOC had lymphatic metastasis and ascites, while patients without SOC did not. The clinical tumour marker cancer antigen 125 (CA125) in two of eight SOC patients was within normal range (<35 U/mL) and only one patient had an abnormal alpha fetoprotein (AFP) level (>7 ng/mL), which reflected the low sensitivity (75% for CA125 and 12.5% for AFP) of the existing clinical protein tumour biomarkers.

We profiled these 16 samples by microarray using the Agilent platform (**Table 1**), yielding comprehensive expression profiles for 17,972 mRNAs, 19,394 lncRNAs, and 5,594 circRNAs. Among these, 3,876 mRNAs, 2,567 lncRNAs, and 191 circRNAs were screened as differentially expressed genes (DEGs) with an absolute fold change >2 and a false discovery rate (FDR) adjusted *P*-value from a Student’s t-test <0.01 (**Figure 3A**). Using GSEA [30], functional enrichment analyses showed that these DEGs are remarkably involved in the Kyoto Encyclopedia of Genes and Genomes (KEGG) [33] pathways related to cancer immunity, including T-cell activation, cytokine production, immune regulation, and tumour-infiltrating lymphocyte differentiation and migration (**Figure 3B and Figure S1**). Therefore, we only concentrated on the immune-related genes for further analysis. According to InnateDB [34] and ImmLnc [35], 884 mRNAs and 426 lncRNAs of these DEGs are immune-related (**Figure 3C and 3D**).

### 3.2 Construction of competing endogenous regulatory network

The competitive endogenous RNA (ceRNA) hypothesis revealed an intrinsic mechanism within RNAs that regulate biological processes and has been validated by many studies [36]. LncRNAs, circRNAs, and mRNAs act as miRNA sponges or ceRNAs by competing for the shared microRNAs (miRNAs). For instance, in the mRNA–miRNA–lncRNA interaction, changes in the expression of lncRNA alter the number of unbound miRNAs, thereby affecting the expression abundance of the target mRNA. To disclose the RNAs participating as ceRNAs in SOC, we screened the positively correlated circRNAs/lncRNAs and mRNAs in SOC samples (Pearson’s correlation coefficient (PCC) >0.5) and selected those that have interactive miRNA in common based on RNAInter [18]. Ultimately, 526 mRNAs, 13 lncRNAs, and 111 circRNAs were identified and composed the competing endogenous regulatory network (**Figure 3E; Additional File 2: Table S1**). Since all the competing endogenous partners are immune-related RNAs, the 111 circRNAs in the ceRNA network were determined as immune-related circRNAs for SOC. The detail description of these immune-related circRNAs is listed in **Additional File 2: Table S3**.

### 3.3 Identification of noncoding RNAs’ competitive endogenous gene pair

In addition to our cohort, we collected all publicly available tissue expression cohorts of SOC from The Cancer Genome Atlas (TCGA) [37] and the Gene Expression Omnibus (GEO) [38], including GSE18520, GSE6008, and GSE40595 (**Table 2; Figure 1B**). We paired two genes if they connected to the same lncRNA or circRNA in the ceRNA regulatory network (**Figure 1A and 3E**) and obtained 186,294 pairs from our self-profiled cohort. Among them, 111,967 pairs existed in the other four public cohorts. We combined the two largest cohorts—TCGA and GSE18520—with our cohort to comprise the training set consisting of 646 patients and 26 normal controls (**Table 2**). The remaining two cohorts served as independent validation sets.

As shown in **Figure 1A**, if two genes that connected to the same noncoding RNA in ceRNA network reversed in their expression between SOC and controls, they were defined as noncoding RNA’s competitive endogenous gene pair (nceGP).

We proposed the denoised individualized pair analysis of gene expression (deiPAGE) (**Figure 2**; see the Methods section) to select nceGPs and constructed a SOC index by extracting and summarising these nceGPs. For each sample in the training cohort, we first calculated the subtraction between two genes regulated by identical ncRNAs in the ceRNA regulatory network (**Figure 2A**). Then, we converted the difference between a pair of genes into a ‘greater’ signal (1) or ‘smaller’ signal (−1) with a noise interval of 0.5 to filter out some false discoveries due to technical variation (**Figure 2B**). If the difference within a pair did not exceed 0.5, the pairing signal was assigned 0. The signals for all possible gene pairs were defined as the relative expression level and we derived a pairwise spectrum of 111,967 pairs. For each pair in the pairwise spectrum, we calculated the contingency table across the population (**Figure 2C**) and 48,388 DEPs were identified (*P* < 0.01, FDR corrected Fisher’s exact test).

Next, we applied LASSO regression model to discriminate the SOC from the controls. After training in the training set, it selected eighteen most effective nceGPs (**Table 3**) to construct the SOC index as follows.

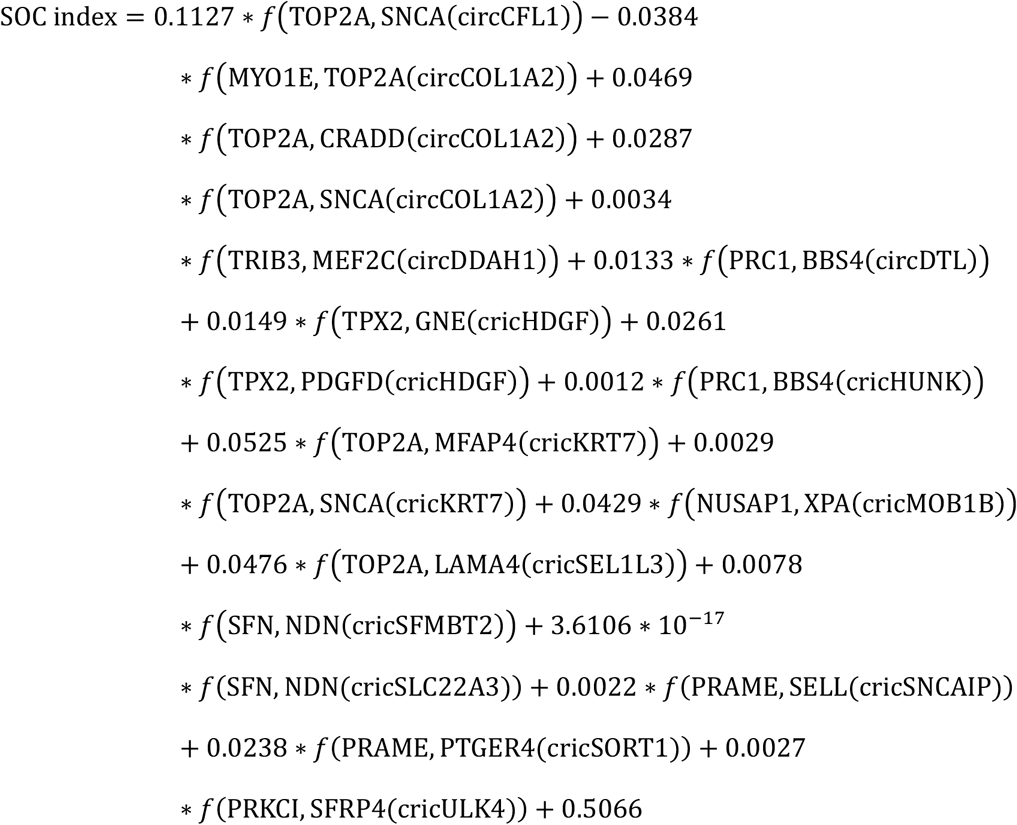

*where* 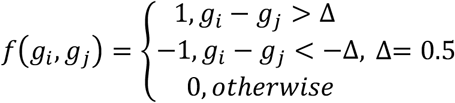.

**Table 3.**
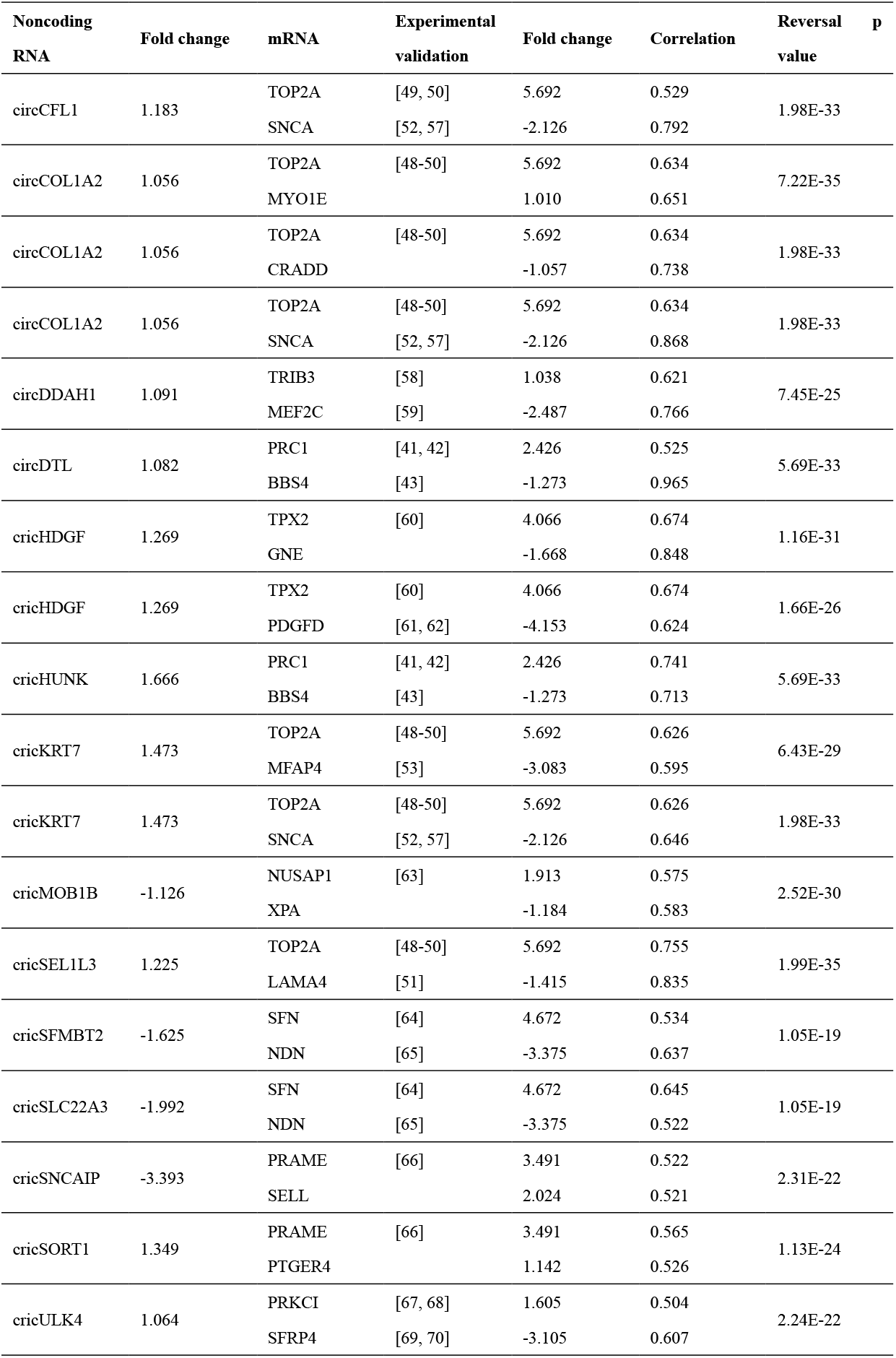
Statistics of the 18 circular RNA’s competitive endogenous gene pairs

Notably, the 18 pairs selected by LASSO were all circRNA’s competing endogenous gene pairs (cceGPs). A heat map displays the effect size (difference between two mRNAs in a pair) of each pair (**Figure 2F**). After conversion, the denoised relative expression shows a clear distinction between SOC patients and normal controls, since all the control samples were clustered in the same group (**Figure 2G**). We further retrieved the 18 cceGPs in ceRNA regulatory network and their common interactive miRNAs (**Figure 3F; Additional File 2: Table S2**). Their locations in the chromosome are shown in circos plot (**Additional File 1: Figure S4**).

Generally, the reversal relationships of the expression abundance of these cceGPs between SOC and control samples were consistent in the training and validation datasets (**Additional File 1: Figure S5, S6 and S7**). Overall, we developed a composite SOC index ranging from 0 to 1.0 on 672 samples in the training set using the LASSO regression model.

### 3.4 Performance of the SOC index

We performed principal components analysis (PCA) of the 18 cceGPs on the training set and two validation sets (**Figure 4A**). The 3D plot of the three principal components illustrates that the 18 pairs selected by deiPAGE revealed differences between the SOCs and controls. The SOC index of SOC samples also significantly differed from controls in the three data sets (**Figure 4B**). Next, we applied the receiver operating characteristic (ROC) curve and the precision-recall curve (PRC) to evaluate the model. The area under the receiver operating characteristic (AUROC) and the area under the precision-recall curve (AUPRC) of SOC index achieved 0.999 and 1.000 in the training set (**Figure 4C**). The SOC index also demonstrated high sensitivity (99.7%) and specificity (92.3%) in discrimination (**Figure 4E**).

To determine if the SOC index obtained from the training set is reproducible in other SOC cohorts, we applied it to two independent validation cohorts (GSE6008 and GSE40595) measured by two different platforms (Affymetrix HG-U133A Array and Affymetrix HG-U133 Plus 2.0 Array). The confusion matrix of GSE6008 (Affymetrix HG-U133A Array) indicated that the SOC index established by deiPAGE from the training set carried a sensitivity of 95.1%, a specificity of 75%, an AUROC of 0.982, and an AUPRC of 0.998 (**Figure 4C, 4D and 4E**: GSE6008). In GSE40595, we assessed the performance of the SOC index to again distinguish SOC from normal using the Affymetrix HG-U133 Plus 2.0 Array platform (**Figure 4C, 4D,** and **4E**: GSE40595). SOC index demonstrated a 100% sensitivity and 100% specificity with an AUROC of 1.000 and an AUPRC of 1.000. Taken together, we observed highly accurate discrimination in the two independent cohorts, indicating that the SOC index carries a very high predictive value in assisting SOC detection, such as improving the accuracy of biopsy diagnosis.

Furthermore, to evaluate the non-invasive diagnosis value of the SOC index, we examined its diagnostic performance in a blood dataset GSE11545. The SOC index achieved comparable AUROC of 0.819 and AUPRC of 0.832 (**Figure 5A and 5B**). Among the 18 cceGPs, we observed that the expression reversal of *PRKCI-cricULK4-SFRP4* and *TOP2A*-circKRT7-*MFAP4* indicated significantly better outcomes in TCGA (**Figure 5C and 5D**).

**Figure 5.**
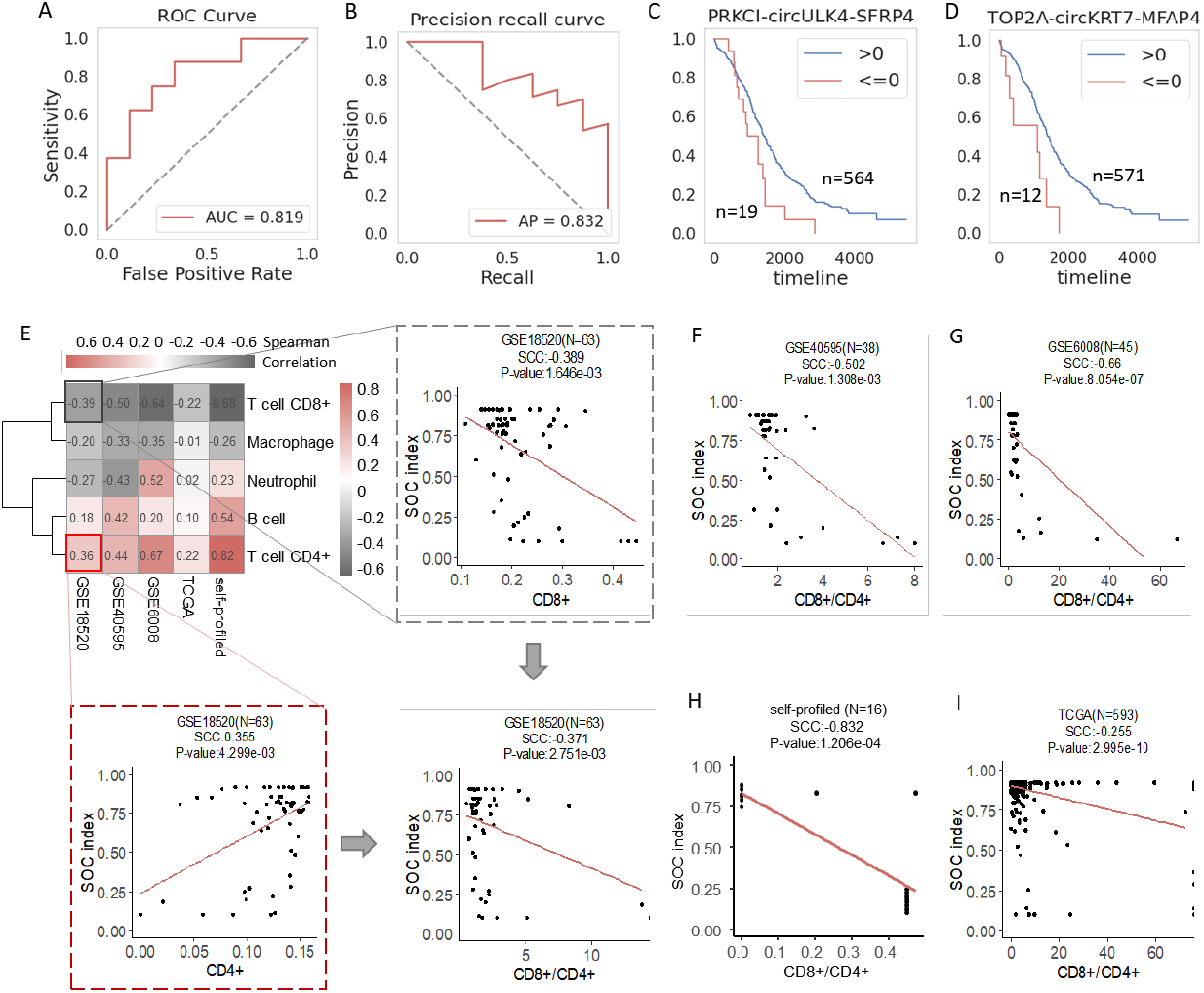
The SOC index for diagnosis and prognosis, and its association with tumor progression. A) ROC curve of the SOC index in blood samples. B) Precision-recall curve of the SOC index in blood samples. C) Kaplan-Meier survival curve of PRKCI:SFRP4 pair (P < 0.03, log-rank test). D) Kaplan-Meier survival curve of TOP2A:MFAP4 pair (P < 0.01, log-rank test). E-I) Correlations between the SOC index and tumour infiltration in five cohorts (E, F, G, H, and I). The SOC index is negatively correlated with CD8+ and positively correlated with CD4+ in GSE18520. The SOC index is also inversely correlated with CD8+/CD4+ (E). In the other four cohorts, the SOC index and CD8+/CD4+ is consistently negatively correlated (F, G, H, and I).

### 3.5 SOC index indicates tumour infiltration

Given that the SOC index of the identified circRNA’s competing endogenous gene pairs potentially discriminated between SOC and normal controls, we questioned whether the score correlated with the tumour progression. We performed a tumour-infiltrating immune cell analysis using TIMER2.0 [39], and decomposed the bulk mRNA expression into cell-type proportions for the self-profiled cohort and the other four cohorts (GSE18520, GSE40595, GSE 6008, and TCGA). We evaluated the correlation between the SOC index and the population of different cell types including CD8+ T cells, CD4+ T cells, B cells, macrophages, and neutrophils on five cohorts (**Figure 5E**). The SOC index was consistently positively associated with CD4+ T cells and negatively associated with CD8+ T cells. For instance in GSE18520, the SOC index was positively correlated with CD4+ T cells (**Figure 5E** and **Additional File 1: Figure S8**; SCC = 0.355, *P* < 4.299E-03, Spearman’s correlation) and negatively correlated with CD8+ T cells (**Figure 5E** and **Additional File 1: Figure S8**; SCC = −0.389, *P* < 1.646E-03), resulting in a significant negative correlation with the ratio of CD8+/CD4+ T cells (SCC = −0.371, *P* < 2.751E-03). The same trend was also observed in the other cohorts (**Figures 5F–I** and **Additional File 1: Figure S8**). The ratio of CD8+/CD4+ T cells in ovarian cancer can be used as a prognostic factor and patients with higher CD8+/CD4+ ratios tend to have improved survival [40]. The high correlation with the CD8+/CD4+ ratio indicates the potential prognostic value of the SOC index.

### 3.6 CircRNA’s competing endogenous gene pairs

The circRNA’s competing endogenous gene pair referred to the correlation between a circRNA and two genes, where the circRNA and two mRNAs are competing endogenous RNA respectively in disease, while the two genes reversed in their expression between the case and control samples. The 18 circRNA’s competing endogenous gene pairs identified by deiPAGE included 14 circRNAs and 22 genes (**Table 3** and **Figure 6A)**. Some gene pairs were regulated by multiple circRNAs while some circRNAs tarteged more than one gene pairs. The 14 circRNAs were differentially expressed with ten up-regulated (circCFL1, circCOL1A2, circDDAH1, circDTL, circHDGF, circHUNK, circKRT7, circSEL1L3, circSORT1, and circULK4) and four down-regulated (circMOB1B, circSFMBT2, circSLC22A3, and circSNCAIP) (**Figure 6B**).

**Figure 6.**
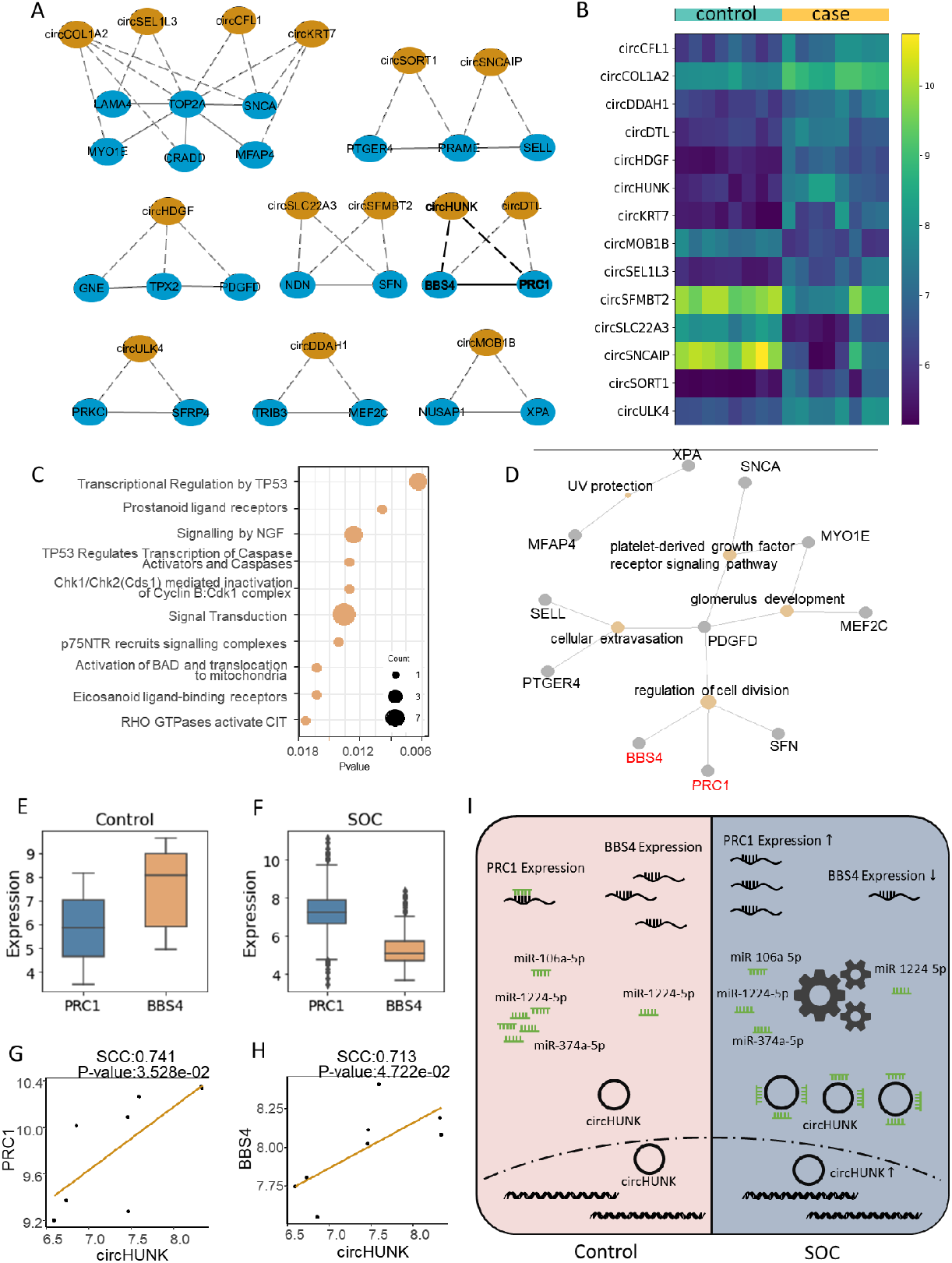
Functional analysis of the cceGPs. A) The 18 cceGPs or gene pair-circRNA motifs in SOC. B) Heatmap illustrating the expression abundance of circRNAs. C) Enriched functions of the 18 cceGPs in Reactome. D) Enriched functions of the 18 cceGPs in GO. Yellow node denotes functional category while grey node represents gene. E, F) Expression abundance of *PRC1* and *BBS4* in normal controls and SOCs. G, H) Correlations between circHUNK and *PRC1 (BBS4*) in SOCs. I) *PRC1*-circHUNK-*BBS4* as an example of cceGPs relationship in SOC progress.

The 22 genes were enriched in Reactome [32] pathways that are highly related to cancer progression, such as signal transduction, signalling by NGF, and transcriptional regulation by TP53 (**Figure 6C**). Other pathways consisted of prostanoid ligand receptors, TP53 regulates transcription of caspase activators and caspases, Chk1/Chk2(Cds1) mediated inactivation of Cyclin B:Cdk1 complex, p75NTR recruits signalling complexes, p75NTR recruits signalling complexes, eicosanoid ligand-binding receptors, and RHO GTPases activate CIT. We also performed functional enrichment in Gene Oncology (GO) [31] and discovered that the genes participated in the cellular growth and development such as regulation of cell division, platelet-derived growth factor receptor signaling pathway, cellular extravasation and glomerulus development (**Figure 6D**). Particularly, Bardet-Biedl Syndrome 4 (*BBS4*) and Protein Regulator Of Cytokinesis 1 (*PRC1*), as competing endogenous RNAs of circHUNK, both participated in the regulation of cell division. Other than that, most of the genes in the 18 circRNA’s competing endogenous gene pairs have been reported to play key roles in carcinogenesis in previous experimental studies (**Table 3**).

Taking *BBS4*–circHUNK–*PRC1* as an example, we dissected the competing endogenous mechanism of the regulatory motif. The expression abundance of *PRC1* was lower than *BBS4* in normal controls, while in SOC samples it is higher than *BBS4* within most of the samples, although both of them are involved in the regulation of cell division in SOC. The boxplot showed the reverse expression pattern of *PRC1* and *BBS4* in the training set (**Figure 6E and 6F**). The expression level of the 18 cceGPs in the training and validation sets were also provided (**Additional File 1: Figure S5, S6 and S7)**. In the SOC samples, circHUNK was significantly positively correlated with *BBS4* and *PRC1* in expression (**Figure 6G** and **6H**). Simultaneously, circHUNK sponged the same microRNA miR-1224-5p as *BBS4* and it shared miR-106a-5p, miR-374a-5p, and miR-1224-5p with *PRC1* (**Figure 6I**). It has been found that the upregulation of *PRC1* activates the Wnt/β-catenin signalling pathway and leads to an increase in cell viability, invasion, migration and EMT of ovarian cancer cells [41]. The overexpression of *PRC1* also indicates a poor prognosis[42]. Moreover, *BBS4* has been observed down-regulated in breast cancer and indicates a shorter survival time [43]. Collectively, these findings suggest that circHUNK play potential roles in the downstream disturbance of cell division in SOV via competing endogenous RNA mechanisms (**Figure 6I**). Our results provide new insight into how the ncRNA-gene pair motif works coordinately in the progression of SOC.

## 4. DISCUSSION

In this study, we hypothesized the triangular relation of mRNA-ncRNA-mRNA that two mRNAs and the ncRNA were competing endogenous RNAs and the two mRNAs were reversed in expression between SOCs and controls. Such mRNA-ncRNA-mRNA motifs were defined as ncRNA’s competing endogenous gene pairs (nceGPs). We constructed a competing endogenous RNA (ceRNA) regulatory network on the basis of the immune-related and differentially expressed RNAs from the self-profiled cohort. We developed deiPAGE algorithm to screen nceGPs and constructed an SOC index for SOC detection, in which 18 nceGPs were included and defined as circRNA’s ceGPs (cceGPs). Validation in two independent cohorts revealed that the SOC index was strongly reproducible and accurate (average AUC close to 0.99).

A preliminary study showed that the six cohorts shared few differentially expressed mRNAs (**Additional File 1: Figure S2 and S3**). For the top 100 DEGs, specifically, only two genes were commonly identified across the six cohorts, indicating that the absolute gene abundance is incapable to consistently identify biomarkers across different platforms. To address this problem, we developed deiPAGE to identify the stable signals from various platforms taking advantage of the relative gene abundance.

The 18 cceGPs consist of 22 genes regulated by 14 circRNAs (**Table 3**). We investigated the 14 circRNAs and found that circKRT7 promotes ovarian cancer cell progression by the circKRT7-miR-29a-3p-*COL1A1* axis [44] and circSLC22A3 suppresses ovarian cancer progression by the CircSLC22A3-miR-518a-5p/Fas axis [45]. CircSFMBT2 inhibits the proliferation and metastasis of glioma cells through miR-182-5p/Mtss1 Pathway [46, 47]. Furthermore, we seek for the experimental studies of the 22 genes and found that 13 genes involve in the regulation in ovarian cancer and three genes participate in the regulation of other cancers. *TOP2A* played a central role in the network as it was involved in four cceGPs (**Figure 6A)**. *TOP2A* (DNA Topoisomerase II Alpha) is a cancer-related gene that encodes DNA topoisomerase enzyme, which controls the structures of DNA during transcription. *TOP2A* promotes tumourigenesis of high-grade SOC by regulating the TGF-β/Smad pathway [48] and serves as the target for several anticancer agents. The *TOP2A* expression is also a marker for the response to pegylated lyposomal doxorubicin (PLD) in epithelial ovarian cancer therapy [49, 50]. In this study, *TOP2A* composed reserved pairs with *LAM4, SNCA, MYO1E, CRADD*, and *MFAP4* in SOC, *LAM4, SNCA*, and *MFAP4* among which are associated with ovarian cancer [51–53]. Particularly, the cceGP *TOP2A:MFAP4* indicates the clinical outcomes of patients (**Figure 5D**). All the genes and circRNAs in cceGPs *TOP2A*-circKRT7-*SNCA* and *TOP2A*-circKRT7-*MFAP4* have been experimental validated to be associated to the regulation of ovarian cancer. Moreover, as *TOP2A*, *SNCA*, and *LAM4* were related to the regulation of ovarian cancer, we speculated that their competing endogenous ncRNAs circCOL1A3 and circSEL1L3 participated in the regulatory mechanism in ovarian cancer.

Function analysis indicated that *BBS4* and *PRC1* in the cceGP *BBS4*-circHUNK-*PRC1* were both enriched in the regulation of cell division (**Figure 6D**). *PRC1* encodes a protein that involves in mitosis and cytokinesis and the overexpression of *PRC1* can infer a poor prognosis in ovarian cancer. Wang et al. found that lncHCP5/miR-525–5p/*PRC1* crosstalk might promote malignant behaviors of ovarian cancer cells and the silencing of lncRNA HCP5 impeded growth and metastasis of tumour in mice [41]. *BBS4* was differentially expressed and downregulated in breast cancer and it was associated with poor prognosis [43]. Collectively, these findings inferred that the circHUNK in this cceGP might be associated with the regulation of gynecologic cancer and might be a potential target for anticancer drugs.

Taking *BBS4*–circHUNK–*PRC1* as an example to dissect the competing endogenous mechanism of the regulatory motif (**Figure 6I**). After the reversed expression of *PRC1* and *BBS4*, the mRNA of *PRC1* turn into a high level and *BBS4* into a low level in SOC. In that case, the mRNAs of *PRC1* and *BBS4* became competing endogenous with circHUNK, as circHUNK sponged common targeted miRNAs with *PRC1* and *BBS4*, and they are positively correlated in SOC. The competing endogenous regulation of the circRNA might be induced by its competing endogenous gene expression reversal.

The proposed SOC index reflects tumour infiltration and is inversely correlated with the ratio of CD8+/CD4+ T-cells, which is lower in the SOC patients and higher among the normal controls. A higher SOC index score implies lower CD8+ fraction and higher CD4+ fraction (**Figures 5E-I**). It suggests that although CD4+ T-cells were active in SOC to help recruit and activate CD8+ T-cells [54], the tumour cells somehow found a way to shut down or deactivate CD8+ cytotoxic T-cells, which can eliminate tumour cells. As the ratio of CD8+/CD4+ T cells in ovarian cancer is used as a prognostic factor [40], the SOC index may be a potential prognostic indicator in cancer patients.

The deiPAGE we proposed in this study is a generalized algorithm for data integration and biomarker identification. Technically, data integration is necessary for a large-scale study to obtain accurate biomarkers [55]. The primary challenge lies on integrating various cohorts, due to the technical variation between platforms and the batch effect from different experiments [56]. To address this issue, the relative expression of gene pairs in each sample rather than the absolute expression value of a single gene was taken into account. Although this might lose some quantitative information from the expression data, datasets from different resources were integrated for model training, thereby substantially increasing the sample size and improving the statistical power of detecting reversal gene pairs. More importantly, the reversal gene pairs can be easily applied to independent individuals, since they do not require any extra preprocessing of population samples. Notably, deiPAGE not only considered the relative ranking but also included the effective size between two genes. Neglecting the size effect difference of two genes for relative expression ordering results in a number of false positives among significant results, given that a large expression difference of two genes contributes equally to a small one. To address this, we used a parameter, difference threshold Δ, to reduce the rate of false-positive discoveries.

Normal ovarian samples are crucial for expression profiling studies relying on comparisons with malignant ovarian tissues. However, tissues from normal donors are rare for ovarian cancer because of the invasive procedure. In total, only 33 normal ovarian samples for gene expression were publicly available among TCGA and GEO before this study. We profiled the expression of eight normal ovarian samples from cervical cancer patients not metastasised to the ovaries, offerring important support and complements to ovarian research. In addition to the mRNA and lncRNA expressions, we measured genome-wide circRNAs using microarray, thereby providing an opportunity to investigate the regulatory role of non-coding RNAs in ovarian cancer. This research may also facilitate studies related to the mechanism and therapeutics involving circRNAs in ovarian cancer.

## 5. CONCLUSION

In this study, we generated a whole transcriptome profile of human ovarian cancer by analysing eight SOCs and eight normal ovary samples using Agilent microarrays, including mRNAs, lncRNAs, and circRNAs. We constructed a competing endogenous RNA (ceRNA) network involving these three types of RNAs and identified immune-related circRNAs from the network. We hypothesized the nceGPs relationship for the mRNA-ncRNA-mRNA motifs in the ceRNA network and proposed deiPAGE to build SOC index and extract 18 cceGPs for SOC. Most of the RNAs in the 18 cceGPs were experimentally validated involved in ovarian cancer development. Our results elucidate the discrimination capability of SOC index and suggest that the competing endogenous motifs play important roles in expression regulation and could be potential target for ovarian cancer mechanism investigation or therapy.

## ABBREVIATIONS

SOC: serous ovarian carcinomas
ceRNA: competing endogenous RNA
deiPAGE: denoised individualized pair analysis of gene expression
ceGPs: competing endogenous gene pairs
nceGPs: noncoding RNA’s competing endogenous gene pairs
cceGPs: cricRNA’s ceGPs
CA125: Cancer antige 125
HE4: human epididymis protein 4
FDA: Food and Drug Administration
PTAR: pro-transition associated R
SNAI2: nail family zinc finger 2
circRNA: circular RNA
ncRNA: long non-coding RNA
GSEA: Gene Set Enrichment Analysis
PCC: Pearson correlation coefficient
DEP: differentially expressed gene pairs
LASSO: least absolute shrinkage and selection operator
TIMER: Tumour Immune Estimation Resource
AFP: bnormal alpha fetoprotein
DEG: differentially expressed genes
KEGG: Kyoto Encyclopedia of Genes and Genomes
miRNA: microRNA
TCGA: The Cancer Genome Atlas
GEO: the Gene Expression Omnibus
DEP: differentially expressed gene pair
AUROC: the area under the receiver operating characteristic
IDH: isocitrate dehydrogenase

## Supplementary Information

**Additional file 1: Figure S1.** The Kyoto Encyclopedia of Genes and Genomes (KEGG) pathways the DEGs involved in besides cancer immunity using Gene Set Enrichment Analysis (GSEA). **Figure S2.** The intersection of top 100 differential up-regulated genes among five cohorts. **Figure S3.** The intersection of top 100 differential down-regulated genes among five cohorts. **Figure S4.** The locations of 18 circRNA’s competing endogenous gene pairs (cceGPs) in the chromosome. **Figure S5.** The expression abundance of the 18 cceGPs were reversed between SOC and control samples in the training dataset. **Figure S6.** The expression abundance of the 18 cceGPs were reversed between SOC and control samples in GSE6008. **Figure S7.** The expression abundance of the 18 cceGPs were reversed between SOC and control samples in GSE40595. **Figure S8.** the correlation between SOC index and CD4+ T cells, CD8+ T cells, the ratio of CD8+/CD4+ T cells in GSE18520, GSE40595, GSE6008, TCGA and self-profiled dataset.

**Additional file 2: Table S1.** mRNAs, circRNAs, lncRNAs in ceRNA Network. **Table S2.** Interactive miRNAs of the 18 ceRNA pairs. **Table S3.** 111 immune-related circRNAs identified by ceRNA network

## DATA AVAILABILITY

The self-profiled data are available at https://www.jianguoyun.com/p/Dbtjn28Qm_jCRjN_ogE.

## ACKNOWLEDGEMENTS

The results are in part based upon data generated by the TCGA Research Network (https://www.cancer.gov/tcga). We are also grateful to Vanessa L Fuller, from Language Services at the University of Helsinki, Finland, for assistance in the English-language revision of this work.

## REFERENCES

1. Guo W, Zhu L, Yu M, Zhu R, Chen Q, Wang Q: A five-DNA methylation signature act as a novel prognostic biomarker in patients with ovarian serous cystadenocarcinoma. Clinical Epigenetics 2018, 10(1):142.

2. Cheng L, Zeng Y, Hu S, Zhang N, Cheung KCP, Li B, Leung KS, Jiang L: Systematic prediction of autophagy-related proteins using Arabidopsis thaliana interactome data. Plant J 2020.

3. Nam EJ, Yoon H, Kim SW, Kim H, Kim YT, Kim JH, Kim JW, Kim S: MicroRNA expression profiles in serous ovarian carcinoma. Clin Cancer Res 2008, 14(9):2690–2695.

4. Hao D, Li J, Wang J, Meng Y, Zhao Z, Zhang C, Miao K, Deng C, Tsang BK, Wang L et al: Non-classical estrogen signaling in ovarian cancer improves chemo-sensitivity and patients outcome. Theranostics 2019, 9(13):3952–3965.

5. Cheng L, Nan C, Kang L, Zhang N, Liu S, Chen H, Hong C, Chen Y, Liang Z, Liu X: Whole blood transcriptomic investigation identifies long non-coding RNAs as regulators in sepsis. J Transl Med 2020, 18(1):217.

6. Sanchez-Mejias A, Tay Y: Competing endogenous RNA networks: tying the essential knots for cancer biology and therapeutics. J Hematol Oncol 2015, 8:30.

7. Chiu HS, Martinez MR, Bansal M, Subramanian A, Golub TR, Yang X, Sumazin P, Califano A: High-throughput validation of ceRNA regulatory networks. BMC Genomics 2017, 18(1):418.

8. Liang H, Yu T, Han Y, Jiang H, Wang C, You T, Zhao X, Shan H, Yang R, Yang L et al: LncRNA PTAR promotes EMT and invasion-metastasis in serous ovarian cancer by competitively binding miR-101-3p to regulate ZEB1 expression. Mol Cancer 2018, 17(1):119.

9. Wang J, Cao Y, Lu X, Wang X, Kong X, Bo C, Li S, Bai M, Jiao Y, Gao H et al: Identification of the Regulatory Role of lncRNA SNHG16 in Myasthenia Gravis by Constructing a Competing Endogenous RNA Network. Mol Ther Nucleic Acids 2020, 19:1123–1133.

10. Cheng L, Leung KS: Quantification of non-coding RNA target localization diversity and its application in cancers. J Mol Cell Biol 2018, 10(2):130–138.

11. Liu X, Xu Y, Wang R, Liu S, Wang J, Luo Y, Leung K-S, Cheng L: A network-based algorithm for the identification of moonlighting noncoding RNAs and its application in sepsis. Briefings in Bioinformatics 2020.

12. Cheng L, Leung KS: Identification and characterization of moonlighting long non-coding RNAs based on RNA and protein interactome. Bioinformatics 2018, 34(20):3519–3528.

13. Wang X, Han L, Zhou L, Wang L, Zhang LM: Prediction of candidate RNA signatures for recurrent ovarian cancer prognosis by the construction of an integrated competing endogenous RNA network. Oncol Rep 2018, 40(5):2659–2673.

14. Li W, Ma S, Bai X, Pan W, Ai L, Tan W: Long noncoding RNA WDFY3-AS2 suppresses tumor progression by acting as a competing endogenous RNA of microRNA-18a in ovarian cancer. Journal of Cellular Physiology 2020, 235(2):1141–1154.

15. Geng Y, Jiang J, Wu C: Function and clinical significance of circRNAs in solid tumors. J Hematol Oncol 2018, 11(1):98.

16. Cheng L, Lo LY, Tang NL, Wang D, Leung KS: CrossNorm: a novel normalization strategy for microarray data in cancers. Sci Rep 2016, 6:18898.

17. Liu X, Li N, Liu S, Wang J, Zhang N, Zheng X, Leung KS, Cheng L: Normalization Methods for the Analysis of Unbalanced Transcriptome Data: A Review. Front Bioeng Biotechnol 2019, 7:358.

18. Lin Y, Liu T, Cui T, Wang Z, Zhang Y, Tan P, Huang Y, Yu J, Wang D: RNAInter in 2020: RNA interactome repository with increased coverage and annotation. Nucleic Acids Res 2020, 48(D1):D189–D197.

19. Nan CC, Zhang N, Cheung KCP, Zhang HD, Li W, Hong CY, Chen HS, Liu XY, Li N, Cheng L: Knockdown of lncRNA MALAT1 Alleviates LPS-Induced Acute Lung Injury via Inhibiting Apoptosis Through the miR-194-5p/FOXP2 Axis. Front Cell Dev Biol 2020, 8:586869.

20. Shannon P, Markiel A, Ozier O, Baliga NS, Wang JT, Ramage D, Amin N, Schwikowski B, Ideker T: Cytoscape: a software environment for integrated models of biomolecular interaction networks. Genome Res 2003, 13(11):2498–2504.

21. Wang J, Xiang X, Bolund L, Zhang X, Cheng L, Luo Y: GNL-Scorer: A generalized model for predicting CRISPR on-target activity by machine learning and featurization. J Mol Cell Biol 2020.

22. Wang J, Zhang X, Cheng L, Luo Y: An overview and metanalysis of machine and deep learning-based CRISPR gRNA design tools. RNA Biol 2020, 17(1):13–22.

23. Tibshirani R: Regression Shrinkage and Selection Via the Lasso. Journal of the Royal Statistical Society: Series B (Methodological) 1996, 58(1):267–288.

24. Liu P, Zheng X, Wong M-H, Leung K-S: Drug2vec: A Drug Embedding Method with Drug-Drug Interaction as the Context. In: International Conference on Engineering Applications of Neural Networks: 2020: Springer; 2020: 326–337.

25. Liu X, Zheng X, Wang J, Zhang N, Leung K-S, Ye X, Cheng L: A long non-coding RNA signature for diagnostic prediction of sepsis upon ICU admission. Clin Transl Med 2020, 10(3):e123.

26. Yang Y, Zhang Y, Li S, Zheng X, Wong MH, Leung KS, Cheng L: A robust and generalizable immune-related signature for sepsis diagnostics. IEEE/ACM Transactions on Computational Biology and Bioinformatics 2021.

27. Li T, Fan J, Wang B, Traugh N, Chen Q, Liu JS, Li B, Liu XS: TIMER: A Web Server for Comprehensive Analysis of Tumor-Infiltrating Immune Cells. Cancer Res 2017, 77(21):e108–e110.

28. Subramanian A, Tamayo P, Mootha VK, Mukherjee S, Ebert BL, Gillette MA, Paulovich A, Pomeroy SL, Golub TR, Lander ES et al: Gene set enrichment analysis: a knowledge-based approach for interpreting genome-wide expression profiles. Proc Natl Acad Sci U S A 2005, 102(43):15545–15550.

29. Yu G, Wang LG, Han Y, He QY: clusterProfiler: an R package for comparing biological themes among gene clusters. OMICS 2012, 16(5):284–287.

30. Subramanian A, Kuehn H, Gould J, Tamayo P, Mesirov JP: GSEA-P: a desktop application for Gene Set Enrichment Analysis. Bioinformatics 2007, 23(23):3251–3253.

31. Gene Ontology C: The Gene Ontology resource: enriching a GOld mine. Nucleic Acids Res 2021, 49(D1):D325–D334.

32. Jassal B, Matthews L, Viteri G, Gong C, Lorente P, Fabregat A, Sidiropoulos K, Cook J, Gillespie M, Haw R et al: The reactome pathway knowledgebase. Nucleic Acids Res 2020, 48(D1):D498–D503.

33. Kanehisa M, Goto S: KEGG: kyoto encyclopedia of genes and genomes. Nucleic Acids Res 2000, 28(1):27–30.

34. Bhattacharya S, Andorf S, Gomes L, Dunn P, Schaefer H, Pontius J, Berger P, Desborough V, Smith T, Campbell J et al: ImmPort: disseminating data to the public for the future of immunology. Immunol Res 2014, 58(2-3):234–239.

35. Li Y, Jiang T, Zhou W, Li J, Li X, Wang Q, Jin X, Yin J, Chen L, Zhang Y et al: Pan-cancer characterization of immune-related lncRNAs identifies potential oncogenic biomarkers. Nat Commun 2020, 11(1):1000.

36. Yang C, Wu D, Gao L, Liu X, Jin Y, Wang D, Wang T, Li X: Competing endogenous RNA networks in human cancer: hypothesis, validation, and perspectives. Oncotarget 2016, 7(12):13479–13490.

37. Cancer Genome Atlas Research N, Weinstein JN, Collisson EA, Mills GB, Shaw KR, Ozenberger BA, Ellrott K, Shmulevich I, Sander C, Stuart JM: The Cancer Genome Atlas Pan-Cancer analysis project. Nat Genet 2013, 45(10):1113–1120.

38. Barrett T, Wilhite SE, Ledoux P, Evangelista C, Kim IF, Tomashevsky M, Marshall KA, Phillippy KH, Sherman PM, Holko M et al: NCBI GEO: archive for functional genomics data sets--update. Nucleic Acids Res 2013, 41(Database issue):D991–995.

39. Li T, Fu J, Zeng Z, Cohen D, Li J, Chen Q, Li B, Liu XS: TIMER2.0 for analysis of tumor-infiltrating immune cells. Nucleic Acids Res 2020, 48(W1):W509–W514.

40. Preston CC, Maurer MJ, Oberg AL, Visscher DW, Kalli KR, Hartmann LC, Goode EL, Knutson KL: The ratios of CD8+ T cells to CD4+CD25+ FOXP3+ and FOXP3-T cells correlate with poor clinical outcome in human serous ovarian cancer. PLoS One 2013, 8(11):e80063.

41. Wang L, He M, Fu L, Jin Y: Role of lncRNAHCP5/microRNA-525-5p/PRC1 crosstalk in the malignant behaviors of ovarian cancer cells. Exp Cell Res 2020, 394(1):112129.

42. Bu H, Li Y, Jin C, Yu H, Wang X, Chen J, Wang Y, Ma Y, Zhang Y, Kong B: Overexpression of PRC1 indicates a poor prognosis in ovarian cancer. Int J Oncol 2020, 56(3):685–696.

43. Zhang H, Suo B, Sun XP, Sun Y-X, Qiao H: Bardet-Biedl Syndrome 4 in Early Diagnosis and Prognosis of Breast Cancer. Indian Journal of Pharmaceutical Sciences 2021:145–152.

44. An Q, Liu T, Wang MY, Yang YJ, Zhang ZD, Lin ZJ, Yang B: circKRT7-miR-29a-3p-COL1A1 Axis Promotes Ovarian Cancer Cell Progression. Onco Targets Ther 2020, 13:8963–8976.

45. Zhang N, Jin Y, Hu Q, Cheng S, Wang C, Yang Z, Wang Y: Circular RNA hsa_circ_0078607 suppresses ovarian cancer progression by regulating miR-518a-5p/Fas signaling pathway. J Ovarian Res 2020, 13(1):64.

46. Sun H, Xi P, Sun Z, Wang Q, Zhu B, Zhou J, Jin H, Zheng W, Tang W, Cao H: Circ-SFMBT2 promotes the proliferation of gastric cancer cells through sponging miR-182-5p to enhance CREB1 expression. Cancer management and research 2018, 10:5725.

47. Zhang S, Qin W, Yang S, Guan N, Sui X, Guo W: Circular RNA SFMBT2 inhibits the proliferation and metastasis of glioma cells through mir-182-5p/Mtss1 pathway. Technology in Cancer Research & Treatment 2020, 19:1533033820945799.

48. Gao Y, Zhao H, Ren M, Chen Q, Li J, Li Z, Yin C, Yue W: TOP2A Promotes Tumorigenesis of High-grade Serous Ovarian Cancer by Regulating the TGF-beta/Smad Pathway. J Cancer 2020, 11(14):4181–4192.

49. Ghisoni E, Maggiorotto F, Borella F, Mittica G, Genta S, Giannone G, Katsaros D, Sciarrillo A, Ferrero A, Sarotto I: TOP2A as marker of response to pegylated lyposomal doxorubicin (PLD) in epithelial ovarian cancers. Journal of ovarian research 2019, 12(1):1–6.

50. Erriquez J, Becco P, Olivero M, Ponzone R, Maggiorotto F, Ferrero A, Scalzo MS, Canuto EM, Sapino A, Verdun di Cantogno L et al: TOP2A gene copy gain predicts response of epithelial ovarian cancers to pegylated liposomal doxorubicin: TOP2A as marker of response to PLD in ovarian cancer. Gynecol Oncol 2015, 138(3):627–633.

51. Liu Y, Xu Y, Ding L, Yu L, Zhang B, Wei D: LncRNA MEG3 suppressed the progression of ovarian cancer via sponging miR-30e-3p and regulating LAMA4 expression. Cancer Cell Int 2020, 20:181.

52. Bruening W, Giasson BI, Klein-Szanto AJ, Lee VM, Trojanowski JQ, Godwin AK: Synucleins are expressed in the majority of breast and ovarian carcinomas and in preneoplastic lesions of the ovary. Cancer 2000, 88(9):2154–2163.

53. Zhao H, Sun Q, Li L, Zhou J, Zhang C, Hu T, Zhou X, Zhang L, Wang B, Li B et al: High Expression Levels of AGGF1 and MFAP4 Predict Primary Platinum-Based Chemoresistance and are Associated with Adverse Prognosis in Patients with Serous Ovarian Cancer. J Cancer 2019, 10(2):397–407.

54. Wang W, Zou W, Liu JR: Tumor-infiltrating T cells in epithelial ovarian cancer: predictors of prognosis and biological basis of immunotherapy. Gynecol Oncol 2018, 151(1):1–3.

55. Liu S, Zhao W, Liu X, Cheng L: Metagenomic analysis of the gut microbiome in atherosclerosis patients identify cross-cohort microbial signatures and potential therapeutic target. FASEB J 2020, 34(11):14166–14181.

56. Zheng X, Wu Q, Wu H, Leung K-S, Wong M-H, Liu X, Cheng L: Evaluating the Consistency of Gene Methylation in Liver Cancer Using Bisulfite Sequencing Data. Frontiers in cell and developmental biology 2021, 9:1022.

57. Fung KM, Rorke LB, Giasson B, Lee VM, Trojanowski JQ: Expression of alpha-, beta-, and gamma-synuclein in glial tumors and medulloblastomas. Acta Neuropathol 2003, 106(2):167–175.

58. Wang S, Wang C, Li X, Hu Y, Gou R, Guo Q, Nie X, Liu J, Zhu L, Lin B: Down-regulation of TRIB3 inhibits the progression of ovarian cancer via MEK/ERK signaling pathway. Cancer Cell Int 2020, 20:418.

59. Kang H, Kim C, Ji E, Ahn S, Jung M, Hong Y, Kim W, Lee EK: The MicroRNA-551a/MEF2C Axis Regulates the Survival and Sphere Formation of Cancer Cells in Response to 5-Fluorouracil. Mol Cells 2019, 42(2):175–182.

60. Ma S, Rong X, Gao F, Yang Y, Wei L: TPX2 promotes cell proliferation and migration via PLK1 in OC. Cancer Biomark 2018, 22(3):443–451.

61. Zhang M, Liu T, Xia B, Yang C, Hou S, Xie W, Lou G: Platelet-derived growth factor D is a prognostic biomarker and is associated with platinum resistance in epithelial ovarian cancer. International Journal of Gynecologic Cancer 2018, 28(2).

62. Yang C, Zhang M, Cai Y, Rong Z, Wang C, Xu Z, Xu H, Song W, Hou Y, Lou G: Platelet-derived growth factor-D expression mediates the effect of differentiated degree on prognosis in epithelial ovarian cancer. J Cell Biochem 2019.

63. Zhang Y, Huang K, Cai H, Chen S, Sun D, Jiang P: The role of nucleolar spindle-associated protein 1 in human ovarian cancer. J Cell Biochem 2020, 121(11):4397–4405.

64. Bryant CS, Kumar S, Chamala S, Shah J, Pal J, Haider M, Seward S, Qazi AM, Morris R, Semaan A et al: Sulforaphane induces cell cycle arrest by protecting RB-E2F-1 complex in epithelial ovarian cancer cells. Mol Cancer 2010, 9:47.

65. Yang H, Das P, Yu Y, Mao W, Wang Y, Baggerly K, Wang Y, Marquez RT, Bedi A, Liu J: NDN is an imprinted tumor suppressor gene that is downregulated in ovarian cancers through genetic and epigenetic mechanisms. Oncotarget 2016, 7(3):3018.

66. Zhang W, Barger CJ, Eng KH, Klinkebiel D, Link PA, Omilian A, Bshara W, Odunsi K, Karpf AR: PRAME expression and promoter hypomethylation in epithelial ovarian cancer. Oncotarget 2016, 7(29):45352–45369.

67. Sarkar S, Bristow CA, Dey P, Rai K, Perets R, Ramirez-Cardenas A, Malasi S, Huang-Hobbs E, Haemmerle M, Wu SY et al: PRKCI promotes immune suppression in ovarian cancer. Genes Dev 2017, 31(11):1109–1121.

68. Rehmani H, Li Y, Li T, Padia R, Calbay O, Jin L, Chen H, Huang S: Addiction to protein kinase Cɩ due to PRKCI gene amplification can be exploited for an aptamer-based targeted therapy in ovarian cancer. Signal transduction and targeted therapy 2020, 5(1):1–11.

69. Belur Nagaraj A, Knarr M, Sekhar S, Connor RS, Joseph P, Kovalenko O, Fleming A, Surti A, Nurmemmedov E, Beltrame L et al: The miR-181a-SFRP4 Axis Regulates Wnt Activation to Drive Stemness and Platinum Resistance in Ovarian Cancer. Cancer Res 2021, 81(8):2044–2055.

70. Ford CE, Jary E, Ma SS, Nixdorf S, Heinzelmann-Schwarz VA, Ward RL: The Wnt gatekeeper SFRP4 modulates EMT, cell migration and downstream Wnt signalling in serous ovarian cancer cells. PLoS One 2013, 8(1):e54362.

