## Supplementary Figures & Tables for "Noncoding RNA’s competing endogenous gene pair as motif in serous ovarian cancer"

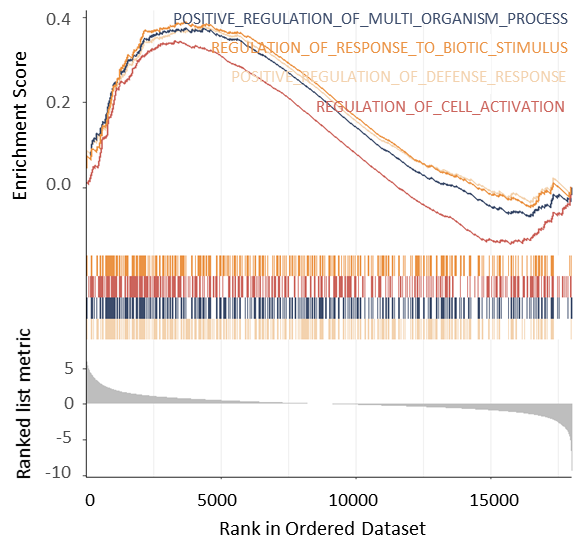

**Figure S1.** Functional analysis of the differentially expressed mRNAs using GSEA. Besides immunity, the enriched biological processes also include some common regulations in cells.

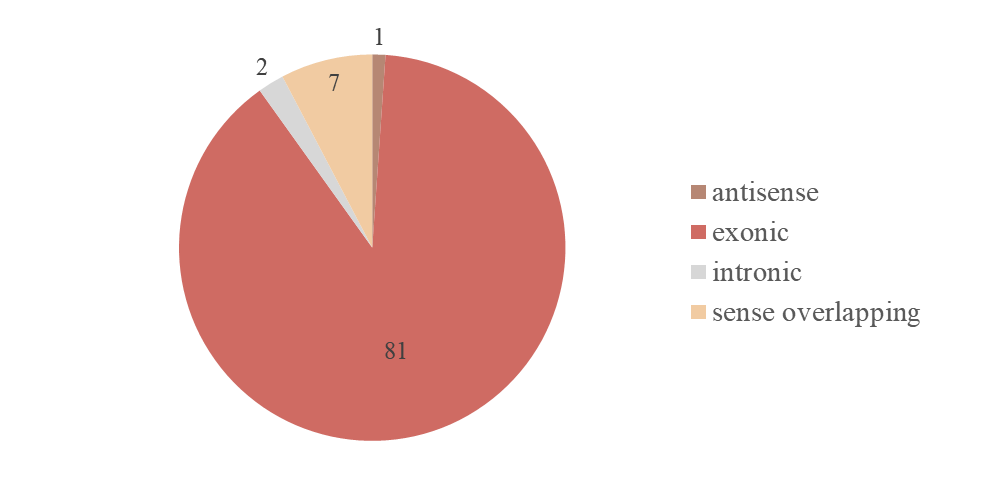

**Figure S2.** CircRNA types of the 91 immune-related circRNAs involved in the competing endogenous regulatory network.

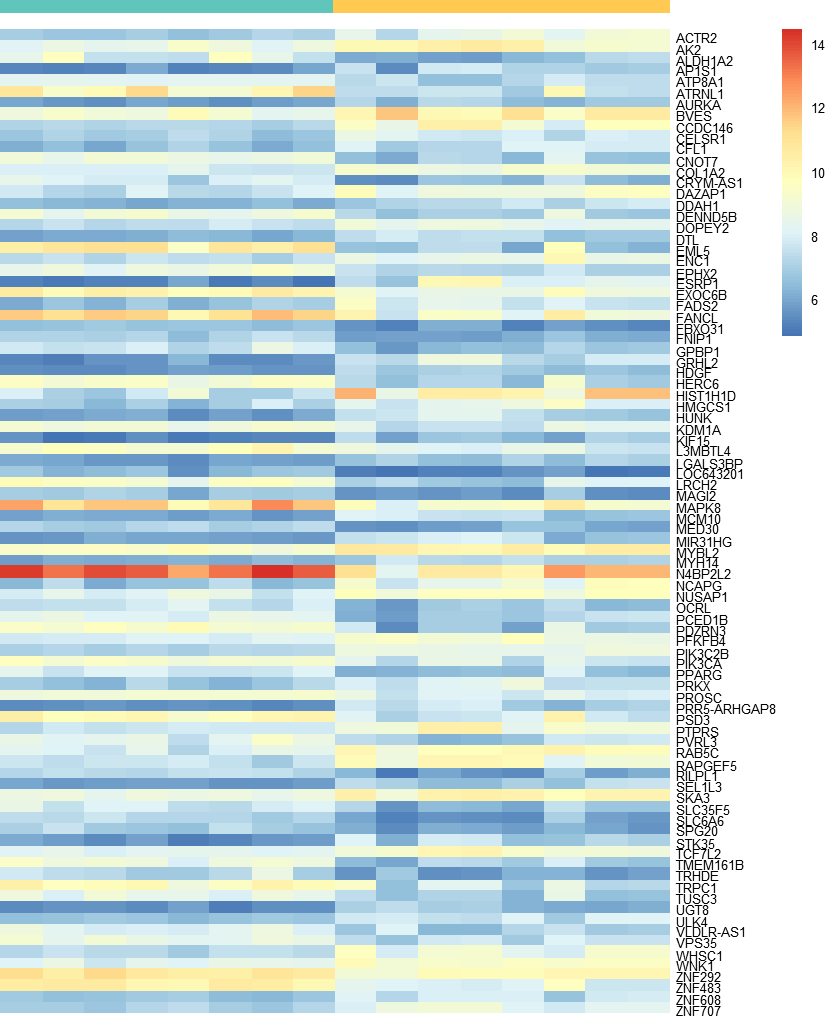

**Figure S3.** Heat map of the 91 immune-related circRNAs involved in the competing endogenous regulatory network. (green bar for normal; yellow bar for disease on the top)

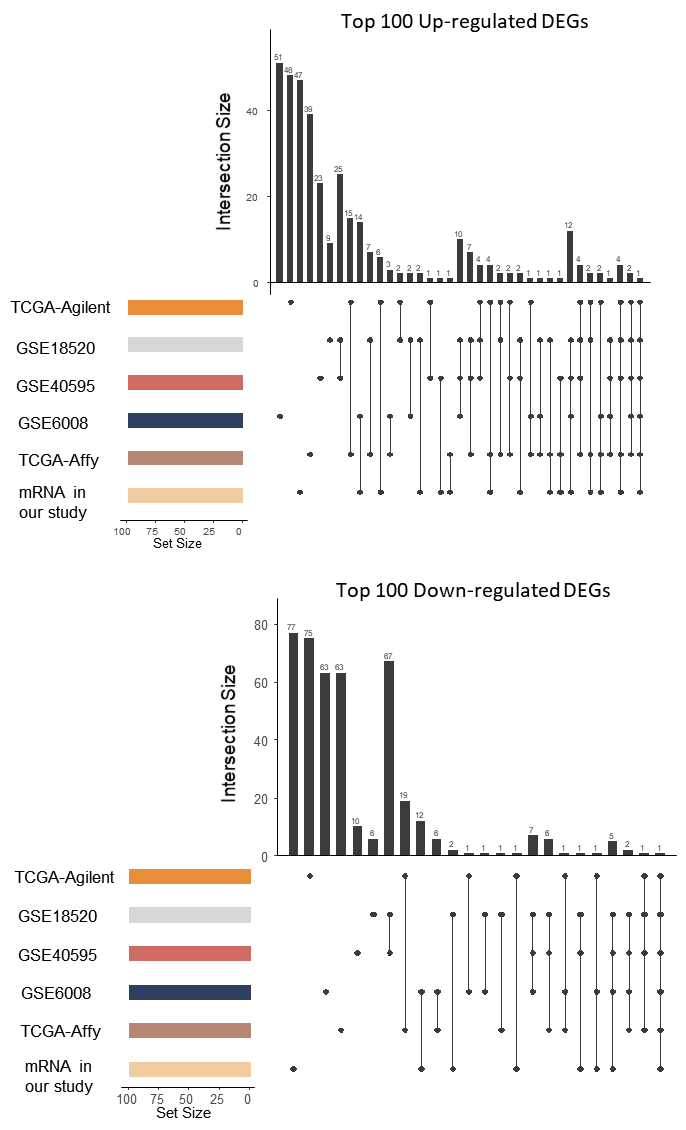

**Figure S4.** The intersection of up-regulated and down-regulated mRNAs among the six ovarian cancer cohorts.

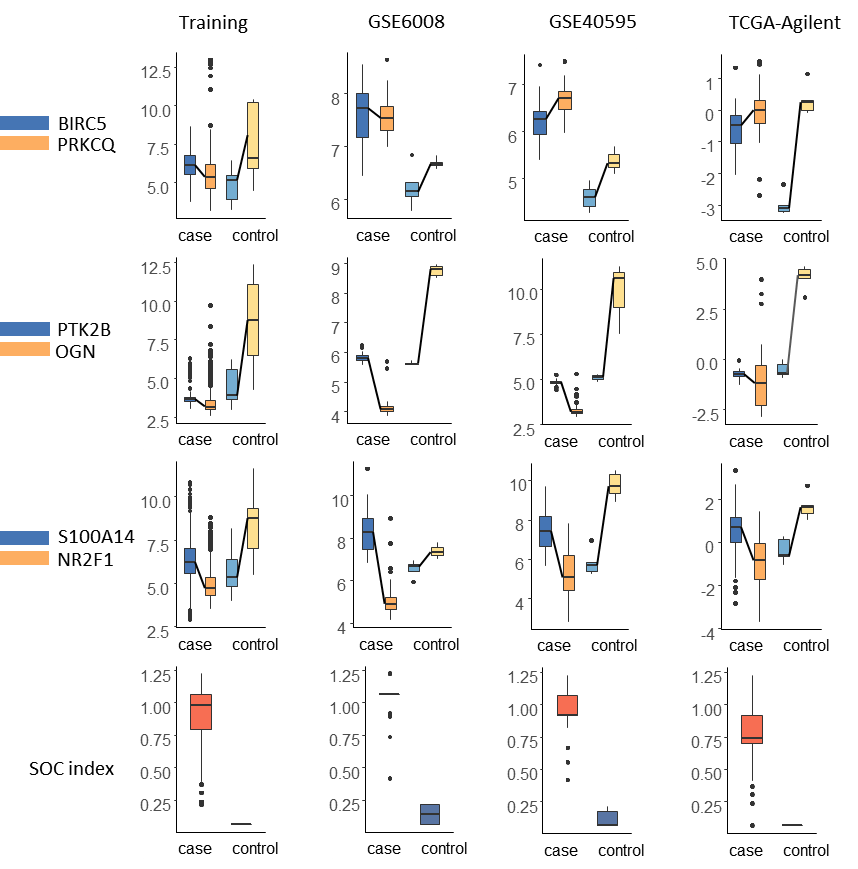

**Figure S5.** Boxplot of the three mRNA pairs and the SOC index in the training set and three validation cohorts.

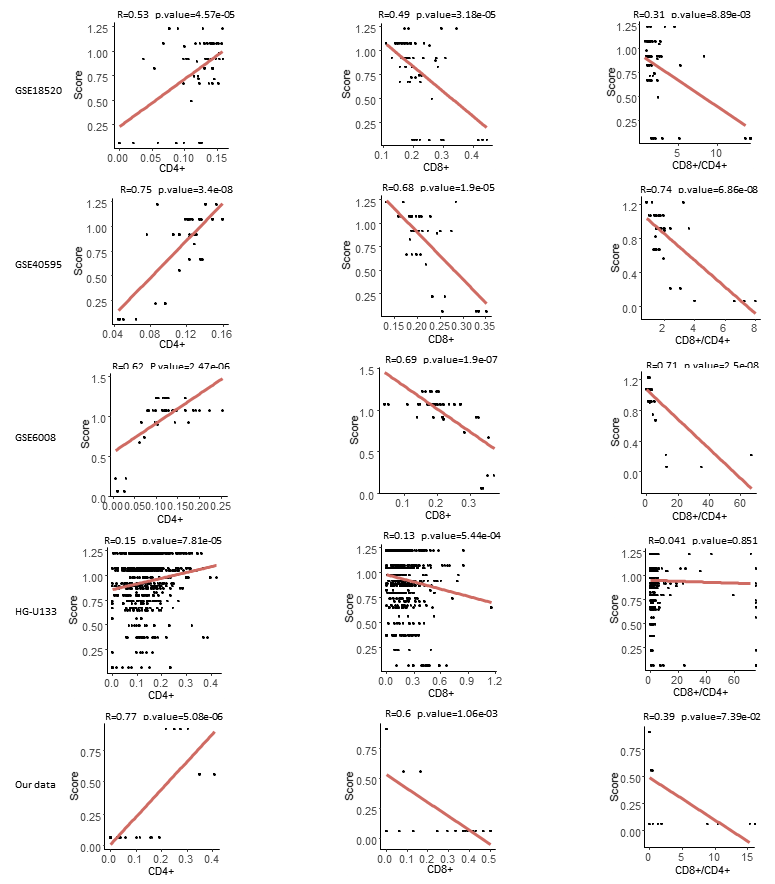

**Figure S6.** Correlation between the SOC index and CD4+ (left column), CD8+ (middle column), as well as CD8+/CD4+ (right column), in the five cohorts.

**Table S1** Clinical and laboratory characteristics of patients with ovarian cancer and control samples.

| **ID** | **Age** | **FIGO staging system** | **Tissue** | **Lymphatic metastasis** | **Ascites** | **AFP** | **CA125** | **Tumour size** | **Tumour size** |
| --- | --- | --- | --- | --- | --- | --- | --- | --- | --- |
|  |  |  |  |  |  | **(ng/mL)** | **(U/mL)** | **(left; cm)** | **(right; cm)** |
| A1 | 53 | Cervical squamous carcinoma stage IB1 | NOET | no | no | – | – | – | – |
| A2 | 52 | Uterine sarcoma | NOET | unknown | unknown | – | – | – | – |
| A3 | 41 | Endometrial carcinoma stage IA | NOET | no | no | – | 522 | – | – |
| A4 | 54 | Cervical squamous cell carcinoma stage IB1 | NOET | no | no | – | – | – | – |
| A5 | 53 | Cervical squamous cell carcinoma stage IB1 | NOET | no | no | – | – | – | – |
| A6 | 61 | Cervical squamous cell carcinoma IIA2 | NOET | unknown | unknown | – | – | – | – |
| A7 | 55 | Cervical squamous carcinoma stage IB1 | NOET | no | no | – | – | – | – |
| A8 | 51 | Endometrial carcinoma stage IA | NOET | no | no | – | – | – | – |
| C1 | 66 | Papillary serous cystadenocarcinomas stage IIIC | SOC | yes | yes | 6.37 | 122.5 | 5*2.5*2.5 | 2.5*2.5*1.2 |
| C2 | 66 | Papillary serous cystadenocarcinomas stage IV | SOC | yes | yes | 3.44 | 15.53 | 8*7*7 | 4*3*3 |
| C3 | 50 | High-grade papillary serous cystadenocarcinomas stage IIIC | SOC | yes | yes | 2.77 | 3239 | 11*10*5 | 5.5*4*2.5 |
| C4 | 51 | Poorly differentiated papillary serous cystadenocarcinomas stage IIIC | SOC | yes | yes | 3.84 | 18.9 | 9*7*4 | 3*1.5*1,5 |
| C5 | 47 | Papillary serous cystadenocarcinomas stage IIIC | SOC | yes | yes | 2.38 | 1818 | 15*14*5 |  |
| C6 | 41 | Ovarian mucinous cystadenocarcinomas stage IIIC | SOC | yes | yes | 8.6 | 115 | 3*2*1 | 4*3*2 |
| C7 | 55 | High-grade papillary serous cystadenocarcinomas stage IIIC | SOC | yes | yes | 2.96 | 730.9 | 1.8*1.3*0.9 | 1.8*1.2*0,7 |
| C8 | 47 | Papillary serous cystadenocarcinomas stage IIIC | SOC | yes | yes | 2.15 | 1278 | 15*2*10 | 3*2.5*2.5 |

Note: FIGO, International Federation of Gynaecology and Obstetrics; NOET, normal ovarian epithelial tissue; AFP, alpha fetoprotein; SOC, serous ovarian carcinomas.

**Table S2** Gene expression datasets used in this study.

| **Dataset** | **Cohorts** | **Platform** | **SOC** | **Control** | **Total** |
| --- | --- | --- | --- | --- | --- |
| Training | TCGA-Affy | Affymetrix HG-U133A Array | 585 | 8 | 593 |
|  | GSE18520 | Affymetrix HG-U133 Plus 2.0 Array | 53 | 10 | 63 |
|  | Own data | Agilent-033010 | 8 | 8 | 16 |
| Test 1 | GSE6008 | Affymetrix HG-U133A Array | 41 | 4 | 45 |
| Test 2 | GSE40595 | Affymetrix HG-U133 Plus 2.0 Array | 32 | 6 | 38 |
| Test 3 | TCGA-Agilent | Agilent G4502A | 32 | 5 | 37 |

**Table S3** Genome characteristics of the three gene pairs.

| **Symbol** | **Chrom** | **Strand** | **Start** | **End** | **EntrezID** | **Description** |
| --- | --- | --- | --- | --- | --- | --- |
| BIRC5 | chr17 | + | 76210276 | 76221716 | 332 | Baculoviral IAP repeat containing 5 |
| PRKCQ | chr10 | - | 6469104 | 6622263 | 5588 | Protein kinase C, theta |
| S100A14 | chr1 | - | 153586731 | 153588808 | 57402 | S100 calcium binding protein A14 |
| NR2F1 | chr5 | + | 92919042 | 92930315 | 7025 | Nuclear receptor subfamily 2 group F member 1 |
| OGN | chr9 | - | 95146248 | 95166937 | 4969 | Osteoglycin |
| PTK2B | chr8 | + | 27179918 | 27316908 | 2185 | Protein tyrosine kinase 2 beta |

| **Table S4**. The 91 immune-related circRNAs involved in the competing endogenous regulatory network. | | | | | | | | | | | |
| --- | --- | --- | --- | --- | --- | --- | --- | --- | --- | --- | --- |
| probeID | circRNA | Alias | GeneSymbol | source | chrom | strand | txStart | txEnd | circRNA_type | best_transcript | Sequence |
| ASCRP3009059 | hsa_circRNA_406521 |  | UGT8 | 25070500 | chr4 | + | 115540578 | 115544858 | sense overlapping | NM_003360 | GCCCACTACCAGAATGAGAGTAACACATGCGCACTGTGGATTCCTTTGTGAATACATAAA |
| ASCRP3004771 | hsa_circRNA_104720 | hsa_circ_0006566 | ZNF707 | circBase | chr8 | + | 144771373 | 144772293 | exonic | NM_173831 | CATGGCCCAGTTGCTGAACCTGTTTGCATGAGTTGCTCCTGACGGCCCTTTAGGATACTT |
| ASCRP3006873 | hsa_circRNA_104651 | hsa_circ_0084927 | ESRP1 | circBase | chr8 | + | 95676924 | 95677424 | exonic | NM_017697 | CTTAAAATTGCTGGTGCAGCAAGATGGAACTTATTGATGATAACACCGTAGTCAGGGCAC |
| ASCRP3000367 | hsa_circRNA_404643 |  | PIK3C2B | 25070500 | chr1 | - | 204396775 | 204397348 | exonic | NM_002646 | AACAATAAACTCTTCATCATGGTGATGCATATTCGGGGCTTGTGTGATTTGGTGTACACC |
| ASCRP3011086 | hsa_circRNA_403102 |  | SEL1L3 | 25242744 | chr4 | - | 25848915 | 25849486 | exonic | NM_015187 | TATTTGGAACCTTCGGGCAAACAGGATTCCACAGTGTCCTCTGGAAAATGAATGTTGTAC |
| ASCRP3004649 | hsa_circRNA_406137 |  | HUNK | 25070500 | chr21 | + | 33331154 | 33347029 | exonic | NM_014586 | CATCTACTTCCTCTTAAACAAGAAACTGGAGCGCTATTTGTCAGGGAGGTGTGAACATGT |
| ASCRP3001445 | hsa_circRNA_406716 |  | LOC643201 | 25070500 | chr5 | - | 175581753 | 175586199 | exonic | NR_036494 | ACGAGATTCAGGAATGACATCAGTACTGGGAAAATGCAAGAAAGTATGTTCCGGTGAAGA |
| ASCRP3005396 | hsa_circRNA_004116 | hsa_circ_0004116 | MED30 | circBase | chr8 | + | 118540889 | 118543066 | exonic | NM_080651 | CCTCGTTTTGCTAGTGAAGAGAGGCGAGAAATTGCTGAAGTAAATAAACTGCCAAATGGT |
| ASCRP3001144 | hsa_circRNA_405028 |  | TRHDE | 25070500 | chr12 | + | 72955944 | 73015531 | exonic | NM_013381 | ACAGGCATCAACACTTATTTCAGATTGGATTTCCAGCAACAGGAACAGGATTATTTAACC |
| ASCRP3012936 | hsa_circRNA_103095 | hsa_circ_0008777 | AURKA | circBase | chr20 | - | 54956488 | 54961589 | exonic | NM_003600 | TTCAGAAACTTTCAAAGTTTGATGAGCAGAGAACTGCTACTGCTACAGCTCCAGTTGGAG |
| ASCRP3011117 | hsa_circRNA_406960 |  | CCDC146 | 25070500 | chr7 | + | 76922268 | 76924255 | sense overlapping | NM_020879 | TCAACCAGGCTTCCTTGTACCCACAGGTGAAAAATATGAATGGCTATCAAAGAAGGATCA |
| ASCRP3002066 | hsa_circRNA_104669 | hsa_circ_0085173 | GRHL2 | circBase | chr8 | + | 102570646 | 102571040 | exonic | NM_024915 | AGCCACAGAGAAACTGCCTTGGCACCAGTGAAGCCCAGAGTAATTTGAGTGGAGGAGAAA |
| ASCRP3003915 | hsa_circRNA_004226 | hsa_circ_0004226 | CDKN2B-AS1 | circBase | chr9 | + | 22046749 | 22066352 | exonic | NR_003529 | CACCATGCATGTGTCCCTTTTGATGAGAAGAATAAGCCTCATTCTGATTCAACAGCAGAG |
| ASCRP3003823 | hsa_circRNA_100684 | hsa_circ_0020048 | TCF7L2 | circBase | chr10 | + | 114724314 | 114799885 | exonic | NM_001146274 | CACACATTGTCCTCCATTTTCAGTCCGGCAGCACACATTACTCTGCGTACAAAACGATTG |
| ASCRP3007855 | hsa_circRNA_401801 |  | RAB5C | 25242744 | chr17 | - | 40278772 | 40282608 | exonic | NM_004583 | GACATCAGCAAAGACTGCAATGAACGTGAACGAAATCTTCATGGCAATAGTTGGAGGTCC |
| ASCRP3003034 | hsa_circRNA_101231 | hsa_circ_0000467 | SKA3 | circBase | chr13 | - | 21742126 | 21742538 | exonic | NM_145061 | ATGCGAGGAATAATAAAAGTACACGAGCAAGAAGCCATTAACTCTGACCCAGAGTTGTCT |
| ASCRP3003129 | hsa_circRNA_100833 | hsa_circ_0022383 | FADS2 | circBase | chr11 | + | 61605249 | 61615756 | exonic | NM_004265 | ATGTGAACATGCTGCACGTGTTTGTTCTGGGCGAATGGCAGCCCATCGAGGATGCCTTCC |
| ASCRP3003784 | hsa_circRNA_405964 |  | CCNT2-AS1 | 25070500 | chr2 | - | 135624111 | 135626611 | sense overlapping | NR_036549 | CAATTCTGGGACAACAAGGGATTCTGGAAGTGTCAGGAAAAGGCAATCACAGGAATATAT |
| ASCRP3012686 | hsa_circRNA_103486 | hsa_circ_0001345 | TRPC1 | circBase | chr3 | + | 142455220 | 142467302 | exonic | NM_003304 | TGTGTTCTGCAAAAAACAAAAAGGATAGCCTCCGGCATTCCAGGTGACTATTATATGGTT |
| ASCRP3013133 | hsa_circRNA_000328 | hsa_circ_0000328 | CFL1 | circBase | chr11 | - | 65622881 | 65623563 | intronic | ENST00000527752 | TTCTTTATAGGGATCAAGCATGAATTGCAAGCAAACTGCTACGAGGAGGGCAAGGAGATC |
| ASCRP3012255 | hsa_circRNA_003300 | hsa_circ_0003300 | LRCH2 | circBase | chrX | - | 114357089 | 114364762 | exonic | NM_020871 | TGCTTGTAAAAAGTTGGGTGTCTCACAGTATAAATCAATGAGGAAGAGTTCAAGTGGCAA |
| ASCRP3006089 | hsa_circRNA_402915 |  | PVRL3 | 25242744 | chr3 | + | 110830876 | 110866331 | exonic | NM_001243288 | CTCAGAAAGACCTATTTCAGGTGCCTTAGCTGGACCAATTATTGTGGAGCCACATGTCAC |
| ASCRP3005000 | hsa_circRNA_403883 |  | AP1S1 | 25242744 | chr7 | + | 100799874 | 100800766 | exonic | NM_001283 | CCACCGATACGTGGAGCTCTTAGACAAATACTTTGGCAGTATGCGGTTCATGCTATTATT |
| ASCRP3012950 | hsa_circRNA_005219 | hsa_circ_0005219 | STK35 | circBase | chr20 | + | 2097311 | 2098061 | exonic | NM_080836 | TTCAGGGCTAAGCATTTTGGGTGATTTTAAACTAGGAGAAAGGATCCTGGGTTATGCTGA |
| ASCRP3011716 | hsa_circRNA_104153 | hsa_circ_0004058 | ZNF292 | circBase | chr6 | + | 87925620 | 87928449 | exonic | NM_015021 | TCTTTCCCAGGAACCATTGGATAAGGATAAAGACACTCCTAGAATATGCAGAGAAATGGA |
| ASCRP3013216 | hsa_circRNA_039210 | hsa_circ_0039210 | VPS35 | circBase | chr16 | - | 46710494 | 46711310 | exonic | NM_018206 | TTGCTTTAATTGATAGATTGTTTTGACTGGCATATTGGAGCAAGTTGTAAACTGTAGGGA |
| ASCRP3004850 | hsa_circRNA_405719 |  | PTPRS | 25070500 | chr19 | - | 5214371 | 5220364 | exonic | NM_002850 | CATTCTGCGTCAGGACATTCTCTCTGCACAAGGAGCTATTTCATTGTGATGGTGCCACTG |
| ASCRP3000220 | hsa_circRNA_407327 |  | OCRL | 25070500 | chrX | + | 128691301 | 128692978 | exonic | NM_000276 | AAAGCGAGAGAAAGAATATGTCAACATTCAGACTTTCAGGTGGCTGCAAAATTCGGGTTC |
| ASCRP3007336 | hsa_circRNA_103066 | hsa_circ_0006332 | MYBL2 | circBase | chr20 | + | 42331129 | 42333998 | exonic | NM_002466 | GAAACATGCTGCGACCCTGATGCTTGGTGTGACCTGAGTAAATTTGACCTCCCTGAGGAA |
| ASCRP3002965 | hsa_circRNA_100295 | hsa_circ_0013451 | SORT1 | circBase | chr1 | - | 109910029 | 109912211 | exonic | NM_002959 | ATGACTTTTGGACAGTCCAAGCTATATCGAAGCATGTGTTTGATGATCTCAGAGGCTCAG |
| ASCRP3012126 | hsa_circRNA_402833 |  | ULK4 | 25242744 | chr3 | - | 41831152 | 41877463 | exonic | NM_017886 | TGCCATGTTGTCCTGTGGGATTCATCTTCAAAGACTAATCCAAGAAAAGGCAATTGTTCT |
| ASCRP3012956 | hsa_circRNA_402760 |  | PRR5-ARHGAP8 | 25242744 | chr22 | + | 45197956 | 45210638 | exonic | NM_181334 | TTCATCAAGGTCCTGTGGAACATCTTGAAGCCCCTCATCAGGGGATGACCGCTTTGGAAG |
| ASCRP3002325 | hsa_circRNA_101529 | hsa_circ_0035442 | ALDH1A2 | circBase | chr15 | - | 58284902 | 58287337 | exonic | NM_003888 | AGGGTCTACTGAGATGGAGACTATTTTACCTTTACAAGACATGAACCCATTGGAGTGTGT |
| ASCRP3010310 | hsa_circRNA_100157 | hsa_circ_0011462 | AK2 | circBase | chr1 | - | 33478807 | 33480195 | exonic | NM_013411 | GTGTTCGCAAGCATCCTAGCAGCCTTCTCCAAAGCCACATGCTGATTCACCCCAAGAGTG |
| ASCRP3007629 | hsa_circRNA_001589 | hsa_circ_0001589 | HIST1H1D | circBase | chr6 | - | 26234499 | 26234709 | sense overlapping | NM_005320 | GAAGGCAGCTCCGAAGAAAAACATCAAAAAGACTCCTAAGAAGGTAAAGAAGCCAGCAAC |
| ASCRP3002887 | hsa_circRNA_103335 | hsa_circ_0006084 | KIF15 | circBase | chr3 | + | 44826336 | 44835918 | exonic | NM_020242 | ATCTGAAAGGCAAAAAGATACCCATGCAGAAGGGATGAGATTGAAGGACCATCTGAATCT |
| ASCRP3007299 | hsa_circRNA_001387 | hsa_circ_0001387 | WHSC1 | circBase | chr4 | + | 1902352 | 1936989 | exonic | NM_007331 | GAAAAGACTCAGGACGGACAAGCACAGTCTTCGGAAGTGTTCTAAGAACGGAAGCATCTG |
| ASCRP3005613 | hsa_circRNA_102747 | hsa_circ_0008529 | ACTR2 | circBase | chr2 | + | 65473657 | 65492309 | exonic | NM_005722 | CGAGTTTTGAAGGGTGATGTGGAAAAACTTTCTGATCTTATGGTTGGTGATGAGGCAAGT |
| ASCRP3009211 | hsa_circRNA_100554 | hsa_circ_0002109 | MCM10 | circBase | chr10 | + | 13214375 | 13214765 | exonic | NM_018518 | GCTCGAACACCAAAGGCTTCACCTCCAGAGGAATTAAGGAATTTGCAAGAGCAAATGAAG |
| ASCRP3009776 | hsa_circRNA_404126 |  | VLDLR-AS1 | 25242744 | chr9 | - | 2536358 | 2539552 | exonic | NR_015375 | GGTCACAAACAAGAGGTCCCTGGATCTACAGGCTCCAAGCTTGATATAGAAGACTATCAT |
| ASCRP3003666 | hsa_circRNA_405649 |  | L3MBTL4 | 25070500 | chr18 | - | 6301901 | 6312055 | exonic | NM_173464 | CAAGGATAGCACAACCCCTTTGAGTCACGATTGCTATGCTATGCCTTGTATGACTAAGAC |
| ASCRP3000535 | hsa_circRNA_034693 | hsa_circ_0034693 | NUSAP1 | circBase | chr15 | + | 41648236 | 41650456 | exonic | NM_016359 | AAGAACACAATTCCATGAATGAACTGAAGGTAACAGAGATTCAAAGGTACCTTCAGAAGG |
| ASCRP3011829 | hsa_circRNA_103927 | hsa_circ_0001523 | ZNF608 | circBase | chr5 | - | 124036706 | 124036962 | exonic | NM_020747 | TGAATTTGGAAGGGATCGTGTGGCATGAAACAGAAGAAGGTTGACCCCCTGTTTACAGTG |
| ASCRP3009693 | hsa_circRNA_103124 | hsa_circ_0001187 | DOPEY2 | circBase | chr21 | + | 37619814 | 37620866 | exonic | NM_005128 | GGGATCCTCTTCCGTTTACTTTAAAACCACCAAACACAACCTCCAAGAGGGAAAACATTT |
| ASCRP3002185 | hsa_circRNA_014784 | hsa_circ_0014784 | HDGF | circBase | chr1 | - | 156711898 | 156713247 | exonic | NM_004494 | AGGAAAATAAAACCCGTTGATTACTATACCTGTAGCCACCAATGTTTCAAGAGGAGCCCC |
| ASCRP3009472 | hsa_circRNA_092549 | hsa_circ_0001486 | GPBP1 | circBase | chr5 | + | 56526672 | 56527148 | exonic | NM_022913 | GGAGGAAAGAAGACAAGAGAGAACGCAAACAGTTTGAAGCTGAGGATTTTTCGTCATTGA |
| ASCRP3009451 | hsa_circRNA_101245 | hsa_circ_0000471 | N4BP2L2 | circBase | chr13 | - | 33091993 | 33101669 | exonic | NM_033111 | GGAGACAGAGAATTCTGCTTGGTCAGAATCGTGATGGCATTGTGTTCAGCACTGATGACT |
| ASCRP3009745 | hsa_circRNA_004607 | hsa_circ_0004607 | DDAH1 | circBase | chr1 | - | 85816097 | 85824530 | exonic | NM_012137 | GGGTCTAGTGAATCTGCACAGAAGGCCCTTAAGGTTGACATGATGAAAGAAGCATTAGAA |
| ASCRP3011578 | hsa_circRNA_103936 | hsa_circ_0073859 | FNIP1 | circBase | chr5 | - | 131006155 | 131014868 | exonic | NM_133372 | TCTCATGCTGTGCAGAACAACAATTTGTAATCTTTACACGATGCCACGAATTGGAGAACC |
| ASCRP3009795 | hsa_circRNA_081069 | hsa_circ_0081069 | COL1A2 | circBase | chr7 | + | 94027693 | 94037203 | exonic | NM_000089 | GGTGAACCTGGTGCCCCTGGTGAAAATGGAACTCCAGGTCAAACAGAAACTGTAAGAAAG |
| ASCRP3009986 | hsa_circRNA_102402 | hsa_circ_0048232 | DAZAP1 | circBase | chr19 | + | 1417498 | 1422395 | exonic | NM_018959 | GTCACGGAGGTAGTCATGATCTATGACGCCGAGAAGCAGAGGCCCCGAGGAAGCTCTTCG |
| ASCRP3005237 | hsa_circRNA_405454 |  | CRYM-AS1 | 25070500 | chr16 | + | 21327227 | 21328447 | sense overlapping | NR_026675 | CCCTCAAATGAAAGAAGAATGGATTTCTCGGAGTCTGAAAAATTTATGGTTCTTCTCTGG |
| ASCRP3007271 | hsa_circRNA_005476 | hsa_circ_0005476 | TMEM161B | circBase | chr5 | - | 87516379 | 87541609 | sense overlapping | NM_153354 | CTTTTGCAATGTATATCTGTCAACAGTATCTCATCATGTTTGGTGATTTGGAGGAAGGGG |
| ASCRP3002759 | hsa_circRNA_405000 |  | PCED1B | 25070500 | chr12 | + | 47606379 | 47610472 | intronic | NM_001281429 | ATCACCAGTTACTTTAATGATCTATTCCTGCAGTGATGGTGTCATACCAGCCCATGGAAC |
| ASCRP3009420 | hsa_circRNA_100097 | hsa_circ_0009061 | KDM1A | circBase | chr1 | + | 23356961 | 23377013 | exonic | NM_015013 | GGACCACAACAGACCCAGAAGGTTTTTCTTTTCATTAGAAACCGCACAGTAGAGTACAGA |
| ASCRP3008327 | hsa_circRNA_101196 | hsa_circ_0007552 | RILPL1 | circBase | chr12 | - | 123983090 | 123984083 | exonic | NM_178314 | GAGGAGGAGCCTGAGGCATGTCAGAGCGGGAGCGACAGGTGATGAAGAAGCTGAAGGAGG |
| ASCRP3001027 | hsa_circRNA_102806 | hsa_circ_0008827 | SLC35F5 | circBase | chr2 | - | 114486977 | 114493435 | exonic | NM_025181 | ACAAGTTGGATATTCCAATGTTCTTTGTGGTTTTTGGCAAATTTGTCATATCAAGAAGCA |
| ASCRP3010478 | hsa_circRNA_100443 | hsa_circ_0000179 | DTL | circBase | chr1 | + | 212218001 | 212220759 | exonic | NM_016448 | GTTTGAGAAAGCTCCCAATATGGAACATGTACTAGCAGTTGCCAATGAAGAAGGCTTTGT |
| ASCRP3013132 | hsa_circRNA_089829 | hsa_circ_0089829 | PRKX | circBase | chrX | - | 3544459 | 3560012 | exonic | NM_005044 | TTCTTGCAGGCAAAATAGATTTCCCCAGACATTTGGATTTCCATGTAAAGACTTGGACCC |
| ASCRP3003409 | hsa_circRNA_104575 | hsa_circ_0083766 | EPHX2 | circBase | chr8 | + | 27382878 | 27394372 | exonic | NM_001979 | CAAAAGCCTCTTCAGAGCAAGCGATGAGGGCGGTGGCCAGTTTGAATACTCCCTTCATAC |
| ASCRP3005632 | hsa_circRNA_102761 | hsa_circ_0009043 | EXOC6B | circBase | chr2 | - | 72945231 | 72960247 | exonic | NM_015189 | CGCAAACATTCAGACAAAATTGGAGAGACTGCCATGAAGCAAAATCAAGTGACGGATACT |
| ASCRP3007816 | hsa_circRNA_102725 | hsa_circ_0004435 | FANCL | circBase | chr2 | - | 58425728 | 58459247 | exonic | NM_018062 | GACACCTCAGGGAAGAGACTTCCACCTTAGGATAGTGTTGCCTGAAGATTTACAACTGAA |
| ASCRP3002110 | hsa_circRNA_018315 | hsa_circ_0018315 | MAPK8 | circBase | chr10 | + | 49609654 | 49613024 | exonic | NM_002750 | AGCGGGCCTACAGAGAGCTAGTTCTTATGAAATGTGTTAATCACAAAAATCTTCTTGGTG |
| ASCRP3006570 | hsa_circRNA_006973 | hsa_circ_0006973 | PDZRN3 | circBase | chr3 | - | 73651504 | 73657835 | exonic | NM_015009 | CTGCAAATTCATGACAGGATTATTGAGGGCGAAGAAACCAAAAGTCTGACTCTTGTCCTG |
| ASCRP3001643 | hsa_circRNA_001788 | hsa_circ_0001788 | PROSC | circBase | chr8 | + | 37623043 | 37623873 | exonic | NM_007198 | ACTTCATTGGCCACCTACAGAAACAAAATGTCAACAAATTGATGGGATCTCCCAGCCATC |
| ASCRP3011212 | hsa_circRNA_001989 | hsa_circ_0001989 | SPG20 | circBase | chr13 | - | 36886281 | 36886614 | exonic | NM_015087 | TCAGATACAATTGATGGAGTTTGCACTGTAGCAAATTGCGTTGGAAAAGAACTAGCTCCA |
| ASCRP3010033 | hsa_circRNA_104166 | hsa_circ_0077527 | BVES | circBase | chr6 | - | 105563560 | 105573453 | exonic | NM_007073 | GTCCTCTCTTCGTAAAGATTGAAAAGGAACTCAGTGGCATGTACCGGCGATTGTTTGAAC |
| ASCRP3012188 | hsa_circRNA_001395 | hsa_circ_0001395 | NCAPG | circBase | chr4 | + | 17816475 | 17816981 | exonic | NM_022346 | TGCTCAGAGAGTAATGCTCCTTCAACAAGGTCTTAATGACAGATCAGCATATGCTACTTT |
| ASCRP3005694 | hsa_circRNA_101894 | hsa_circ_0040766 | FBXO31 | circBase | chr16 | - | 87376482 | 87380856 | exonic | NM_024735 | CGGGAGGCAGGAGTGCTTCACCGATATAGACACATTTTGGGATTGTGGCAGCCAGATATC |
| ASCRP3001894 | hsa_circRNA_403022 |  | PIK3CA | 25242744 | chr3 | + | 178921331 | 178922376 | exonic | NM_006218 | AGAGTACCTTGTTCCAATCCCAGTATATAAGAAGCTGTATAATGCTTGGGAGGATGCCCA |
| ASCRP3001831 | hsa_circRNA_001681 | hsa_circ_0001681 | RAPGEF5 | circBase | chr7 | - | 22330793 | 22357656 | exonic | NM_012294 | AGACTCCGAAGAAAGCAGTGATGAAATTCTTGTGCGTCTAACATCTGCGACTCTATCAAG |
| ASCRP3004746 | hsa_circRNA_102592 | hsa_circ_0052001 | MYH14 | circBase | chr19 | + | 50720871 | 50721028 | exonic | NM_024729 | CATGCTGCAGGACGTACTCCGGCCTTTTCTGTGTGGTCATCAACCCGTACAAGCAGCTTC |
| ASCRP3005100 | hsa_circRNA_406276 |  | PFKFB4 | 25070500 | chr3 | - | 48585964 | 48587667 | exonic | uc003ctw.3 | CCATCCACCGCTAAAAAGTGTCCATCATTCCAATATTCCTCAATAGTGTGCATGACCAAC |
| ASCRP3012746 | hsa_circRNA_103631 | hsa_circ_0069618 | ATP8A1 | circBase | chr4 | - | 42553214 | 42558057 | exonic | NM_006095 | TGTTTTCACTGGAAGAACACCCGACTCGGTGATTATAGATTCAGCAGAACTCACAGTTTG |
| ASCRP3000301 | hsa_circRNA_003251 | hsa_circ_0003251 | WNK1 | circBase | chr12 | + | 1003727 | 1006847 | exonic | NM_014823 | AAATCTAGTCGAAGCAGTTCCTTGGGGAATAAAAGCCCCCAGCTTTCAGTTTCTCAAGTC |
| ASCRP3005531 | hsa_circRNA_000810 | hsa_circ_0000810 | LGALS3BP | circBase | chr17 | - | 76967343 | 76968212 | sense overlapping | NM_005567 | CCTTGTCACTGGTCTGAGGTCATTAAAATTACATTGAGGATACCTACAAGCCCCGGATTT |
| ASCRP3013414 | hsa_circRNA_073009 | hsa_circ_0073009 | ENC1 | circBase | chr5 | - | 73923233 | 73925817 | exonic | ENST00000302351 | TTTCTCTTCACTCACTCCATCACTCGATGAGATAATATGAGATTTCTACTTCGGAGAGGC |
| ASCRP3010785 | hsa_circRNA_103830 | hsa_circ_0072389 | HMGCS1 | circBase | chr5 | - | 43294157 | 43299077 | exonic | NM_002130 | TACTCTCTTAAAGTCACACAAGATGCTACACCGGCTCTTTCACCATGCCTGGATCACTTC |
| ASCRP3011858 | hsa_circRNA_100686 | hsa_circ_0020093 | ATRNL1 | circBase | chr10 | + | 116879948 | 116889297 | exonic | NM_207303 | TGACCTGACTGGAGAAAAATTATGTGTCTGCAATGATAGTTGGCAAGGTTAACAGAACCT |
| ASCRP3001879 | hsa_circRNA_104559 | hsa_circ_0008752 | CNOT7 | circBase | chr8 | - | 17092224 | 17094882 | exonic | NM_013354 | AGAGCTGCAAAAATCTCAAAGGAGGACATGTATGCCCAGGACTCTATAGAGCTACTAACA |
| ASCRP3007790 | hsa_circRNA_101039 | hsa_circ_0025821 | DENND5B | circBase | chr12 | - | 31586088 | 31595859 | exonic | NM_144973 | GACAACGACTTGAGGGAGCCCAATCAATTGAGCAGAGACTGATGAAAATGGATCACACTG |
| ASCRP3003151 | hsa_circRNA_101424 | hsa_circ_0032883 | EML5 | circBase | chr14 | - | 89220855 | 89221015 | exonic | NM_183387 | ATGTTCATACACATGGTATAGCTTGCTTGGCGTTTGACTTAGATGGACAGCCTTGCATTG |
| ASCRP3005176 | hsa_circRNA_001304 | hsa_circ_0001425 | HERC6 | circBase | chr4 | - | 89340453 | 89341580 | antisense | NM_017912 | CCCAGTGACTTCTGGATACCACTGACTCTAGATTCATTTCACTACCAGGTCAAAGACACA |
| ASCRP3002230 | hsa_circRNA_403848 |  | MAGI2 | 25242744 | chr7 | - | 77885259 | 77885898 | exonic | NM_012301 | AGGACTGTCCCATTGGAAGTGAAACTTCTTTGATTATCCATCGAGGAGGTGATGTCATTG |
| ASCRP3009289 | hsa_circRNA_064335 | hsa_circ_0064335 | PPARG | circBase | chr3 | + | 12353878 | 12422990 | exonic | NM_138711 | GCTTCTGGATTTCACTATGGAGTTCATGCTTGTGAAGGATGCAAGATTTGAAAGAAGCCA |
| ASCRP3006806 | hsa_circRNA_104566 | hsa_circ_0004458 | PSD3 | circBase | chr8 | - | 18656804 | 18662408 | exonic | NM_015310 | CTGTGCAATAATGCTTCTTAATACCGATCTACATGGCCACCAATGCTGGGGTGAAAACAA |
| ASCRP3002512 | hsa_circRNA_103298 | hsa_circ_0004705 | SLC6A6 | circBase | chr3 | + | 14485131 | 14489324 | exonic | NM_003043 | AGTTCTGGGAAAAGCAAGGAGATGGCCACCAAGGAGAAGCTGCAGTGTCTGAAAGATTTC |
| ASCRP3011259 | hsa_circRNA_104557 | hsa_circ_0006410 | TUSC3 | circBase | chr8 | + | 15508205 | 15531345 | exonic | NM_006765 | TCATAAGAACCCACACAATGGACAAGTGGCAAGCTAATGAAGAATATCAAATACTGGCGA |
| ASCRP3012279 | hsa_circRNA_404246 |  | ZNF483 | 25242744 | chr9 | + | 114293154 | 114296633 | exonic | NM_133464 | TGGAAGAAGTCTCAAAAAGCTCCCGACTAGATCCAGTCTCTCAAGATTCTACTGTTTCCC |
| ASCRP3003382 | hsa_circRNA_008599 | hsa_circ_0008599 | MIR31HG | circBase | chr9 | - | 21476897 | 21477291 | exonic | NR_027054 | ATTTTGTGACTACACTGCAGGTTTCTGGTCCTCATACCGTGTGGTTGAGTGAAATCAAAA |
| ASCRP3004400 | hsa_circRNA_103258 | hsa_circ_0063822 | CELSR1 | circBase | chr22 | - | 46829289 | 46832186 | exonic | NM_014246 | CAGGTGCAGTACTACAACAAGGTTTGCCACTCAGGAAAGGAACGGCTTGCTTCTCTACAA |
